# An activating mutation in *Pdgfrb* causes skeletal stem cell defects with osteopenia and overgrowth in mice

**DOI:** 10.1101/2021.01.21.427619

**Authors:** Hae Ryong Kwon, Jang H. Kim, John P. Woods, Lorin E. Olson

## Abstract

Autosomal dominant PDGFR*β* gain-of-function mutations in mice and humans cause a spectrum of wasting and overgrowth disorders afflicting the skeleton and other connective tissues, but the cellular origin of these disorders remains unknown. We demonstrate that skeletal stem cells (SSCs) isolated from mice with a gain-of-function D849V point mutation in PDGFR*β* exhibit SSC colony formation defects that parallel the wasting or overgrowth phenotypes of the mice. Single-cell RNA transcriptomics with the SSC colonies demonstrates alterations in osteoblast and chondrocyte precursors caused by PDGFR*β*^D849V^. Mutant SSC colonies undergo poor osteogenesis *in vitro* and mice with PDGFR*β*^D849V^ exhibit osteopenia. Increased expression of *Sox9* and other chondrogenic markers occurs in SSC colonies from mice with PDGFR*β*^D849V^. Increased STAT5 phosphorylation and overexpression of *Igf1* and *Socs2* in PDGFR*β*^D849V^ SSCs suggests that overgrowth in mice involves PDGFR*β*^D849V^ activating the STAT5-IGF1 axis locally in the skeleton. Our study establishes that PDGFR*β*^D849V^ causes osteopenic skeletal phenotypes that are associated with intrinsic changes in SSCs, promoting chondrogenesis over osteogenesis.

## Introduction

Two platelet-derived growth factor (PDGF) receptors (PDGFRs) have been identified in mammals, PDGFR*α* and PDGFR*β*, which bind to five PDGF ligands. PDGFRs play crucial and distinct roles in embryo development by regulating the proliferation, migration, survival, and differentiation of mesenchymal cells that populate all tissues and organs (Hoch and Soriano 2003; Andrae et al. 2008; Klinkhammer et al. 2018). It has recently been discovered that humans with gain-of-function mutations in *PDGFRB* exhibit a spectrum of phenotypes affecting the skeleton and other connective tissues in an autosomal-dominant fashion (Guerit et al. 2021). These mutations result in constitutive PDGFR*β* signaling, causing Penttinen syndrome (MIM 601812) or Kosaki overgrowth syndrome (MIM 616592). Both disorders progressively affect the skeleton beginning in childhood. Other activating variants of *PDGFRB* are associated with a milder phenotype, infantile myofibromas (MIM 228550), which does not affect the skeleton. Penttinen syndrome, with *PDGFRB* mutations V665A or N666S (mutated in the first kinase domain), is characterized as a premature aging condition with osteoporosis, scoliosis, lipoatrophy, dermal atrophy, aneurysms, and acro-osteolysis (Johnston et al. 2015; Bredrup et al. 2019). Kosaki overgrowth syndrome, with *PDGFRB* mutations P584R or W566R (mutated in the juxtamembrane domain), is featured by tall stature, elongated long bones, enlarged hands and feet, distintive facial features, scoliosis, hyperelastic skin, aneurysms, myofibromas, and neurodegeneration (Takenouchi et al. 2015; Minatogawa et al. 2017). The pathological mechanisms of the human disorders are still unknown.

We previously demonstrated that mice with a gain-of-function D849V mutation in *Pdgfrb* (mutated in the second kinase domain, corresponding to the human D850 residue) died between 2-3 weeks of age after developing a postnatal wasting phenotype (Olson and Soriano 2011; He et al. 2017). Surprisingly, these phenotypes were modulated by signal transducer and activator of transcription 1 (STAT1). Mice with *Pdgfrb^D849V^* mutation but lacking *Stat1* (*Pdgfrb^+/D849V^Stat1^-/-^*mice) survived until 8-9 weeks while becoming overweight with widespread connective tissue overgrowth (not obesity), thick calvarias, abnormally curved spine, and enlarged rib cage (He et al. 2017). Although the murine model clearly identified *Stat1* as a gene modulating the phenotype spectrum driven by *Pdgfrb^D849V^*, many questions remain about the target cell types and signaling pathways underlying such striking phenotypes. Moreover, the role of PDGFR*β* in the skeleton is not well understood, as PDGFR*β* seems to be redundant for skeletal development based on the normal skeletal phenotypes of global and osteoblast-specific knockouts in mice (Soriano 1994; Bohm et al. 2019).

Ligand binding to wild type PDGFR*β* induces receptor dimerization, which activates the receptor’s kinase activity and results in autophosphorylation of intracellular tyrosine residues that activate downstream signaling pathways (Lemmon and Schlessinger 2010). Gain-of-function PDGFR*β* mutations disrupt the inactive conformation of the receptor, leading to constitutive kinase activity and autophosphorylation. PDGFRs utilize a variety of signaling pathways to mediate their effects on cell behavior, including PI3K, MAPK, PLC*γ*, and STAT1/3/5 (Heldin and Westermark 1999; Tallquist and Kazlauskas 2004; Demoulin and Essaghir 2014). PDGFR*β* is particularly important for pericyte development and function, but it is also expressed on fibroblasts, osteoblasts, and stem/progenitor cells with potential to differentiate into multiple mesenchymal cell types (Andrae et al. 2008).

Skeletal stem cells (SSCs) residing in bone and bone marrow are responsible for postnatal bone development, tissue homeostasis, and repair (Bianco and Robey 2015). A single SSC at the apex of skeletal lineages can give rise to chondrocytes, osteoblasts, adipocytes and fibroblasts. Recent findings with *in vivo* lineage tracing and single-cell transplantation have increased the rigor of SSC biology and improved our understanding of SSC heterogeneity (Ambrosi et al. 2019; Serowoky et al. 2020). Perisinusoidal vasculature in bone marrow (BM) is surrounded by SSCs expressing PDGFR*β*, PDGFR*α*, CD146, Nestin, LepR and Cxcl12, while arterial vasculature is associated with SSCs expressing PDGFR*β*, PDGFR*α*, Sca1 and LepR (Sacchetti et al. 2007; Morikawa et al. 2009; Mendez-Ferrer et al. 2010; Zhou et al. 2014). The resting zone of the growth plate harbors SSCs expressing Grem1, Col2a1, PthrP and Itgav (CD45^-^/TER119^-^/Tie2^-^/Thy^-^/6C3^-^/CD105^-^/CD200^-^/Itgav^+^) (Chan et al. 2015; Worthley et al. 2015; Mizuhashi et al. 2018; Newton et al. 2019). The periosteum and cranial sutures contain SSCs expressing PDGFR*β*, Gli1, Axin2, Ctsk, and *α*SMA (Yang et al. 2013; Zhao et al. 2015; Shi et al. 2017; Debnath et al. 2018; Ortinau et al. 2019). PDGFR*β* is broadly expressed in SSCs, but its role has not been examined. We speculated that PDGFR*β* signaling could be functional in SSCs, and hypothesized that elevated *Pdgfrb^D849V^* mutant signaling in SSCs could alter stem cell functions to generate skeletal disorders.

## Results

### PDGFRβ^D849V^ alters skeletal growth in mice

To explore the PDGFR*β*^D849V^ skeleton, we generated a cohort of mice and measured their weights and bone lengths. As *Stat1* is an important modifier of PDGFR*β*^D849V^ phenotypes (He et al. 2017), four offspring genotypes were established with the *Sox2Cre* driver for germline activation of *Pdgfrb^D849V^* and/or deletion of *Stat1*-floxed alleles by crossing *Pdgfrb^+/D849V^Stat1^flox/flox^* and *Stat1^+/-^Sox2Cre^+/-^* mice. The resulting four genotypes are: *Stat1^+/-^* (hereafter designated as *S^+/-^*), *Stat1^-/-^* (*S^-/-^*), *Pdgfrb^+/D849V^Stat1^+/-^* (*KS^+/-^*), and *Pdgfrb^+/D849V^Stat1^-/-^* (*KS^-/-^*). The *Pdgfrb* allele expressing D849V is designated K because the mutation is in the kinase domain. We used two genetic controls, *S^+/-^* and *S^-/-^*, that do not show growth or survival defects (Meraz et al. 1996; He et al. 2017). *KS^+/-^* mice died by 3 weeks of age with features of autoinflammation and wasting. *KS^-/-^* mice, however, were rescued in survival at 3 weeks and died around 8-9 weeks, consistent with previous findings (He et al. 2017). Thus, body weights were collected from live mice at 2-9 weeks of age, and bone length data were collected from tibias at 3 or 6-8 weeks of age. In the resulting growth curve, *KS^+/-^* and *KS^-/-^* mice were clearly distinguishable by their obvious wasting and overgrowth phenotypes, respectively (Figure 1A). 3-week-old *KS^+/-^* tibias were shorter than *KS^-/-^* and control tibias (Figure 1B), and by 6-8 weeks the *KS^-/-^* tibias were significantly longer than controls (Figure 1C). We also examined osteoclasts by tartrate-resistant acid phosphate (TRAP) staining in each genotype to test whether altered bone resorbing cells could be coupled with the skeleton growth defects. TRAP stain was stronger in both mutants at 3 weeks old (Supplementary Figure 1), which indicates increased osteoclastogenesis in both *KS^+/-^* and *KS^-/-^* mice and does not correlate with the wasting or overgrowth phenotypes. We conclude that changes in osteoclast activity do not explain the skeleton growth phenotypes. Instead, skeletal lineages expressing PDGFR*β* are likely to be responsible for the skeletal growth defects. Therefore, we next investigated the effects of PDGFR*β*^D849V^ on SSCs.

**Figure 1.**
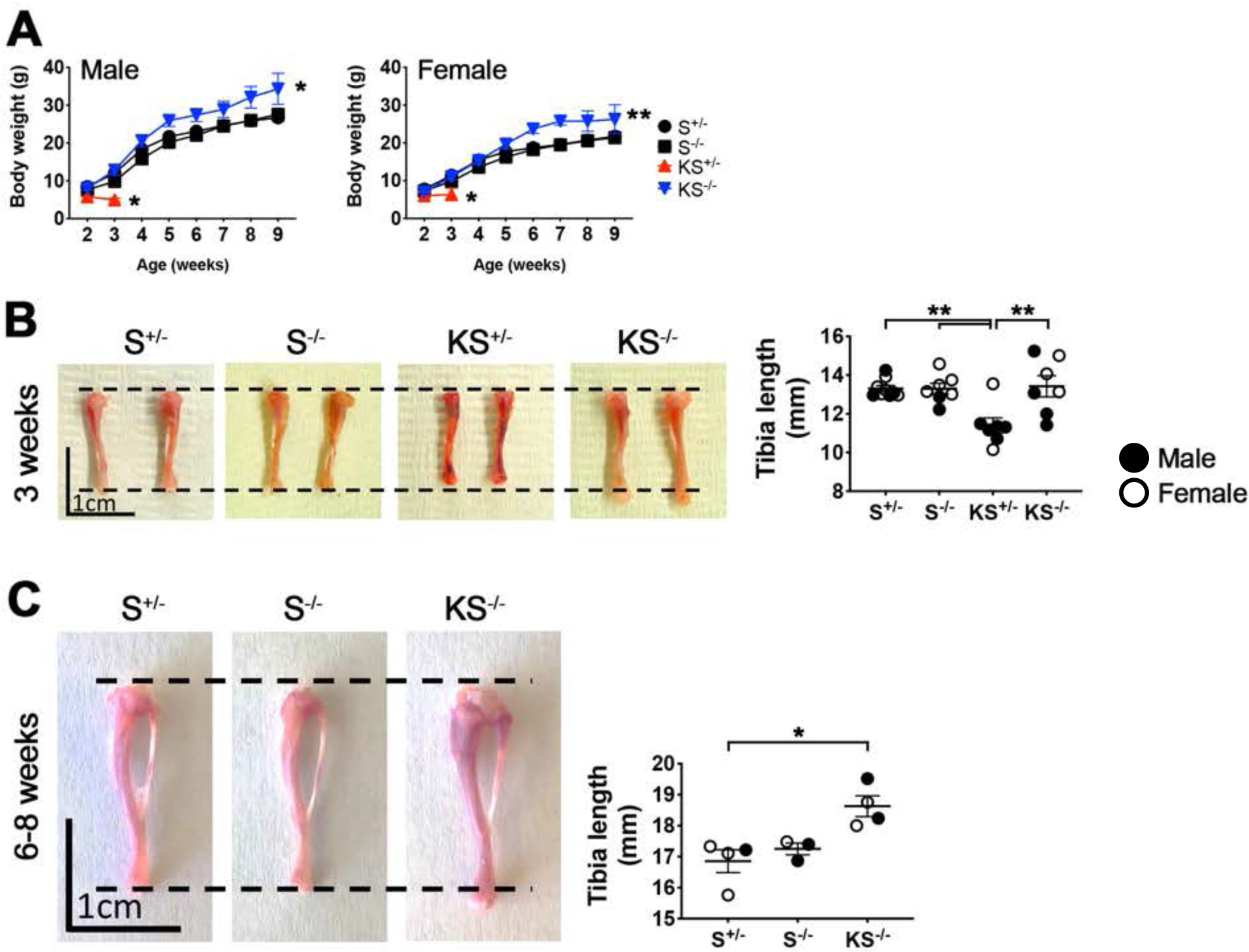
PDGFR*β*^D849V^ alters skeletal growth rate in mice. **(A)** Male and female body weights of 2 to 9-week-old mice of the indicated genotypes (males: n = 5 for *S^+/-^*, 8 for *S^-/-^*, 3 for *KS^+/-^* and 10 for *KS^-/-^*, females: n = 8 for *S^+/-^* and *S^-/-^*, 3 for *KS^+/-^* and 10 for *KS^-/-^*). **, *p* < 0.05 *KS^-/-^* vs *S^-/-^* for 2 to 9-week-old females. *, *p* < 0.05 *KS^-/-^* vs *S^-/-^* for 2 to 9-week-old males or *KS^+/-^* vs *KS^-/-^* for 2 to 3-week-old males and females. **(B)** Tibia length measurements at 3 weeks old (n = 9 for *S^+/-^* and 7 for *S^-/-^*, *KS^+/-^* and *KS^-/-^*). **(C)** Tibia length measurements at 6-8 weeks old (n = 4 for *S^+/-^*, 3 for *S^-/-^* and 4 for *KS^-/-^*). Male and female data were plotted with black and white circles, respectively. Statistics analysis were performed by two-way ANOVA (A) or one-way ANOVA (B and C). Data represent mean ± SEM.

### PDGFRβ^D849V^ regulates SSC numbers and colony formation

To determine whether SSCs were altered in PDGFR*β*^D849V^-expressing mice, we performed SSC quantification and colony formation assays. SSCs were immunophenotyped from limb bones with enzymatic digestion and flow cytometry using two well-established SSC markers, PDGFR*α* and Sca1/Ly6a (P*α*S) (Morikawa et al. 2009; Houlihan et al. 2012) (Supplementary Figure 2). The percentage of P*α*S cells in the stromal fraction (excluding hematopoietic and endothelial cells) was calculated at 3 and 6 weeks of age. The P*α*S percentage was similar between all genotypes at 3 weeks old (Figure 2A), but it was significantly increased in cells isolated from *KS^-/-^* bones at 6 weeks old (Figure 2B). To characterize stem cell function, we sorted out P*α*S cells from 3-week-old and 6 to 9-week-old bones and performed colony formation assays. Western blotting confirmed constitutive activation of PDGFR*β* and knockout of STAT1 in SSC-derived colonies (Figure 2C). At 3 weeks there was a decrease in the number of colonies generated by *KS^+/-^* SSCs compared to equal numbers of *S^+/-^* or *S^-/-^* colonies, while the number of colonies was increased in *KS^-/-^* SSCs (Figure 2D and 2E). The colonies were classified into groups based on size (small (=<5 um), medium (5<x<36), large (=>36)) to examine the expansion capacity of control and mutant SSCs. *KS^+/-^* SSCs generated decreased colony numbers of all three sizes, and *KS^-/-^* increased medium size colonies (Figure 2E). 6 to 9-week-old *KS^-/-^* SSCs also increased the number of colonies formed compared to controls (Figure 2F-2G). These results show that mice with PDGFR*β*^D849V^ exhibit changes in the number of SSCs and colony forming unit activity in parallel to the wasting and overgrowth phenotypes displayed *in vivo*, consistent with the idea of intrinsic defects in mutant SSCs mediating skeletal phenotypes.

**Figure 2.**
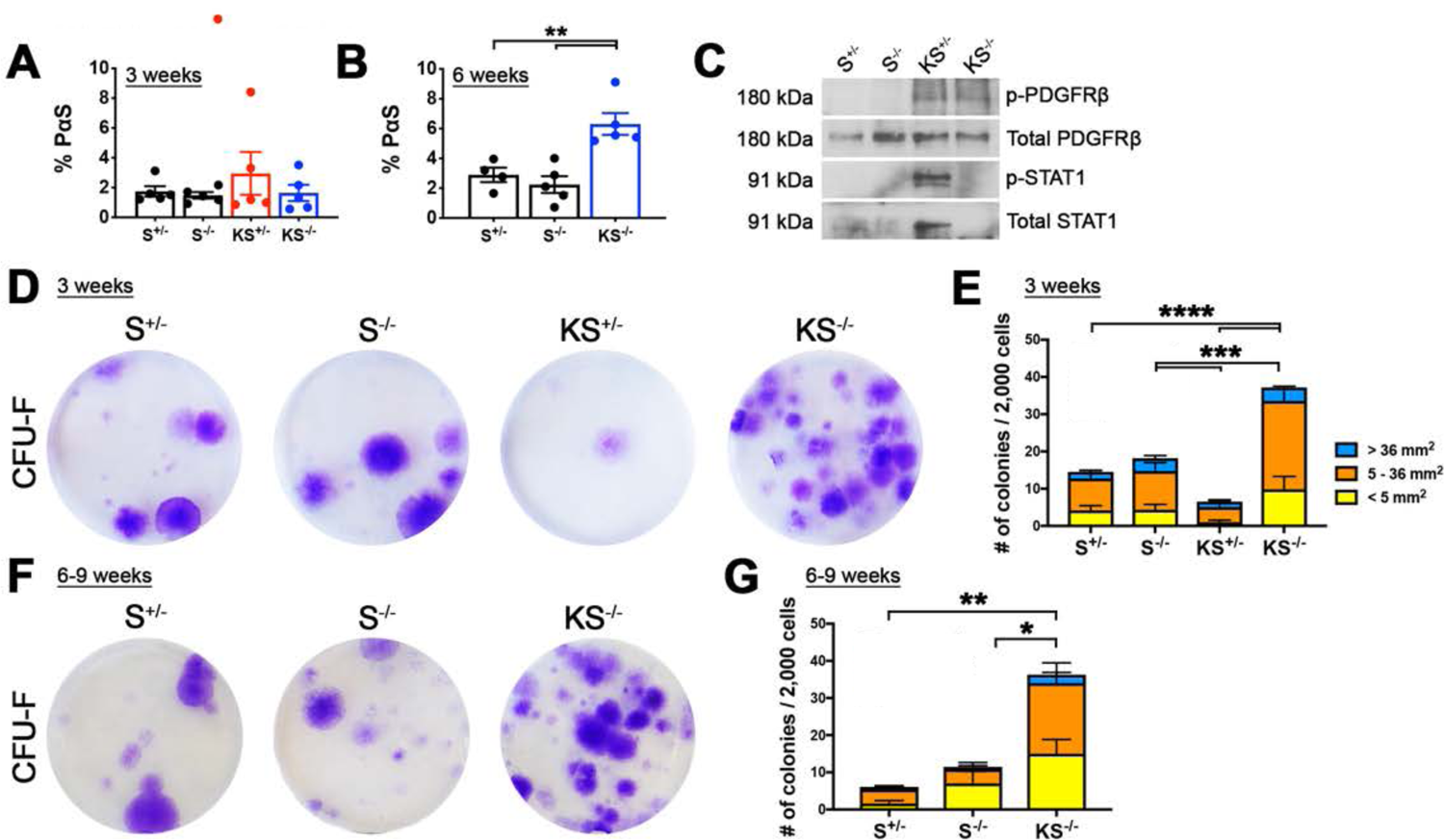
PDGFR*β*^D849V^ alters skeletal stem cell numbers *in vivo* and colony formation capacity *in vitro*. **(A)** Ratio of PDGFR*α*/Sca1 double positive skeletal stem cells (P*α*S SSCs) among stromal fraction (non-hematopoietic and non-endothelial skeletal pool) within 3-week-old long bones (n= 5 for all four genotypes). **(B)** P*α*S SSCs ratio at 6 weeks old (n = 4 for *S^+/-^* and 5 for *S^-/-^* and *KS^-/-^*). **(C)** Phosphorylated PDGFR*β* (pY1009), total PDGFR*β*, phosphorylated STAT1 (pY701), and total STAT1 in SSCs after overnight serum-starvation, representative of two biological replicates. **(D-G)** CFU-F colonies stained with crystal violet and quantified. P*α*S SSCs were isolated by FACS from 3-week-old (D and E) or 6-9-week-old mice (F and G). CFU-F assays were performed with 2,000 P*α*S SSCs isolated from individual mice in a 6-well plate. Quantification plots show total colony numbers and three categories of colony size (n = 6 mice for *S^+/-^*, *S^-/-^* and *KS^-/-^* and 4 for *KS^+/-^* for 3 weeks, n = 5 for *S^+/-^* and *S^-/-^* and 4 for *KS^-/-^* for 6-9 weeks). Statistical analysis of E and G were performed with total numbers of colonies between genotypes. ****, *p* < 0.001, ***, *p* < 0.005, **, *p* < 0.01 and *, *p* < 0.05 by one-way ANOVA. Data represent mean ± SEM.

### Single-cell RNA sequencing indicates multi-lineage potential of cultured SSCs

P*α*S cell-derived colonies can differentiate into osteoblasts or adipocytes when treated with differentiation cocktails (Morikawa et al. 2009). However, we noted that the colonies were only partially differentiated (Supplementary Figure 3), suggesting cellular heterogeneity within each SSC-derived colony. As an approach to evaluate cellular heterogeneity of SSCs and obtain detailed information on single-cell differentiation potential, we performed single-cell transcriptomics. To escape the autoinflammatory condition in *KS^+/-^* mice, which would strongly influence gene expression, we utilized SSC colonies that had been cultured for 14 days without differentiation cocktails instead of freshly isolated cells. Thus, single-cell suspensions of colonies from the four genotypes were subjected to single-cell RNA (scRNA) sequencing using 10X Genomics Chronium platform. We integrated scRNA sequencing data from eight samples, representing the four genotypes in duplicate, to generate general clusters using the Seurat package (Butler et al. 2018; Stuart et al. 2019). Each cluster was grouped based on signature genes that were differentially expressed between clusters (Figure 3A). A heatmap of signature gene marker expression is shown in Figure 3B (full list in Supplementary Table 1). These markers, combined with current literature and gene ontology, were used to define the cell type represented by each cluster (Figure 3C and 3D, Supplementary Table 2). Clusters of biological duplicates were distributed similarly within each genotype (Supplementary Figure 4A). As shown by uniform manifold approximation and projection (UMAP) plot (Figure 3A), we found 10 clusters containing either SSCs, intermediate skeletal stem and progenitor cells (SSPCs), chondrocyte precursors, osteoblast precursors, or adipocyte precursors. All 10 clusters were conserved across the four genotypes (Supplementary Figure 4B).

**Figure 3.**
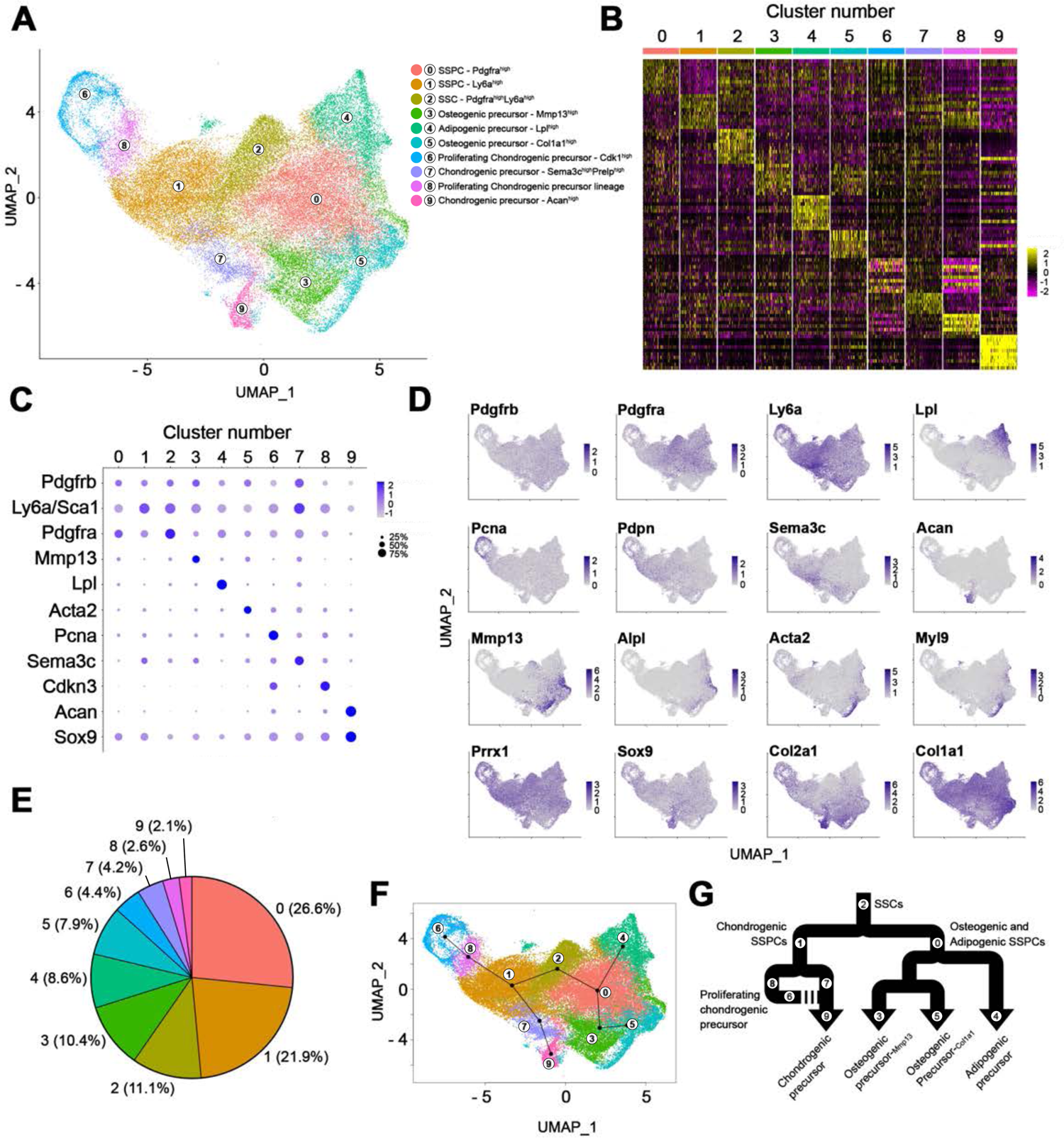
Single-cell transcriptomics reveals multilineage potential of SSC-derived CFU-F colonies. **(A)** UMAP plot showing 10 clusters and their defined cell types. scRNA data from all four genotypes with biological duplicates were integrated and clustered by Seurat. **(B)** Heatmap generated with top markers of 10 clusters (see Supplementary Table 1 for full list). **(C)** Dotplot of highly expressed markers in each cluster. Dot color indicates expression level and dot size indicates percentage of cells. **(D)** UMAP plot highlighting distribution of 16 markers expressed on SSCs, SSPCs and precursors. **(E)** Proportion of each cluster in the total population. **(F)** Pseudo-time trajectory analyzed by Slingshot with all 10 clusters. **(G)** A summary of SSCs to four precursor lineages through intermediate SSPCs based on pseudotime trajectory analysis.

As a percentage of all cells, SSCs and SSPCs were more abundant than the committed precursors (Figure 3E). The most abundant cluster was cluster 0 (26.6%, SSPCs with *Pdgfra^high^*), followed by cluster 1 (21.9%, SSPCs with Sca1/Ly6a^high^) and cluster 2 (11.1%, SSCs with *Pdgfra^high^* and *Sca1/Ly6a^high^*). Cluster 2 was considered the origin of all populations due to high expression of *Pdgfra* and *Sca1/Ly6a*. The mesoderm marker *Prrx1* was most highly expressed in cluster 2 and was broadly expressed in other clusters, as expected for SSCs and their progeny (Supplementary Figure 4C). *Pdgfrb* was moderately expressed in most clusters including SSCs (cluster 2), but was downregulated in chondrocyte precursors (clusters 6, 8 and 9) (Figure 3C). The SSC and SSPC clusters broadly expressed several previously identified stem cell markers including *Prg4*, *Cxcl12*, *Ctsk*, *Cd164*, *Cd51/Itgav* and *Grem1*, but others were barely detected including *Nes*, *Cd146/Mcam*, *Lepr*, and *Gli1* (Supplementary Figure 4C).

The remaining clusters, 3 through 9, were considered precursors rather than differentiated cells, because the colonies were cultured with maintenance medium to support clonal expansion without differentiation. Representative markers for each precursor cluster are summarized in Supplementary Figure 4D. Of note, cluster 3 represents osteogenic precursors expressing *Mmp13*, *Alpl* and *Sp7*. Cluster 4 represents adipocyte precursors expressing *Lpl*, *Fabp4*, *Hp* and *Adipoq*. Cluster 5 represents osteoblastic precursors highly expressing *Col1a1* and *Col1a2* and moderately expressing *Acta2*, *Tagln* and *Myl9*. Clusters 6 and 8 represent actively proliferating chondrocyte precursors highly expressing cell cycle genes (*Cdk1*, *Mki67* and *Pcna*) and early chondrogenic markers (*Col2a1*, *Pdpn* and *Grem1*). Since we regressed out cell cycle genes (S and G2/M phase genes) during dimensional reduction of scRNA data, many S phase cells were distributed throughout all the clusters (Supplementary Figure 4E). However, clusters 6 and 8 remained prominent for cell cycle, chromosome and mitosis gene signatures (Supplementary Table 2). Cluster 7 represents chondrogenic precursors with expression of Sema3c and Prelp. Cluster 9 represents chondrocyte precursors with expression of *Acan*, *Ucma*, *Col9a1*, *Sox9* and *Col2a1*.

Given the initial seeding of P*α*S SSCs (cluster 2) and subsequent emergence of precursors with chondrogenic (clusters 6-9), osteogenic (clusters 3 and 5) and adipogenic (cluster 4) properties, we hypothesize that precursors were generated through intermediate SSPCs (clusters 0 and 1). Pseudotime projection analysis with Slingshot was performed to identify possible branching events representing cell lineages (Street et al. 2018). This suggested two major lineage trajectories from cluster 2 as the top of the hierarchy: one trajectory leads through *Sca1/Ly6a^high^* SSPCs (cluster 1) to chondrogenic precursors, and the other leads through *Pdgfra^high^* SSPCs to osteogenic/adipogenic precursors (Figure 3F). The lineage scheme from SSCs to SSPCs to precursors is summarized in Figure 3G.

### PDGFRβ^D849V^ impairs osteogenesis

To identify changes in clusters and gene expression due to PDGFR*β*^D849V^, we split the scRNA data into controls (*S^+/-^* and *S^-/-^*) and mutants (*KS^+/-^* and *KS^-/-^*). *Pdgfrb* mRNA was moderately decreased in *KS*^+/-^ and *KS*^-/-^ genotypes, as shown previously in dermal fibroblasts (He et al. 2017), and *Stat1* mRNA was absent from *S^-/-^* and *KS^-/-^* (Supplementary Figure 5A). We quantified the abundance of each cluster as a percentage of all cells represented by control or mutant genotypes (Figure 4A). Clusters that were increased or decreased in mutant colonies compared to controls were color-coded in the lineage map summaries (Figure 4B). PDGFR*β*^D849V^ colonies particularly decreased osteogenic clusters (cluster 3 and 5). To identify specific gene expression changes, we further analyzed scRNA data with the Database for Annotation, Visualization and Integrated Discovery (DAVID) analysis (Huang da et al. 2009) (Supplementary Table 3 and 4). We found that PDGFR*β*^D849V^ colonies downregulated osteogenesis-related genes including *Col1a1*, *Col1a2*, *Mmp13*, *Ptx3*, *Serpine1* and *Serpinf1* (Figure 4C). Their expression levels were decreased in almost all clusters, suggesting a broad impact across skeletal lineages (Figure 4D).

**Figure 4.**
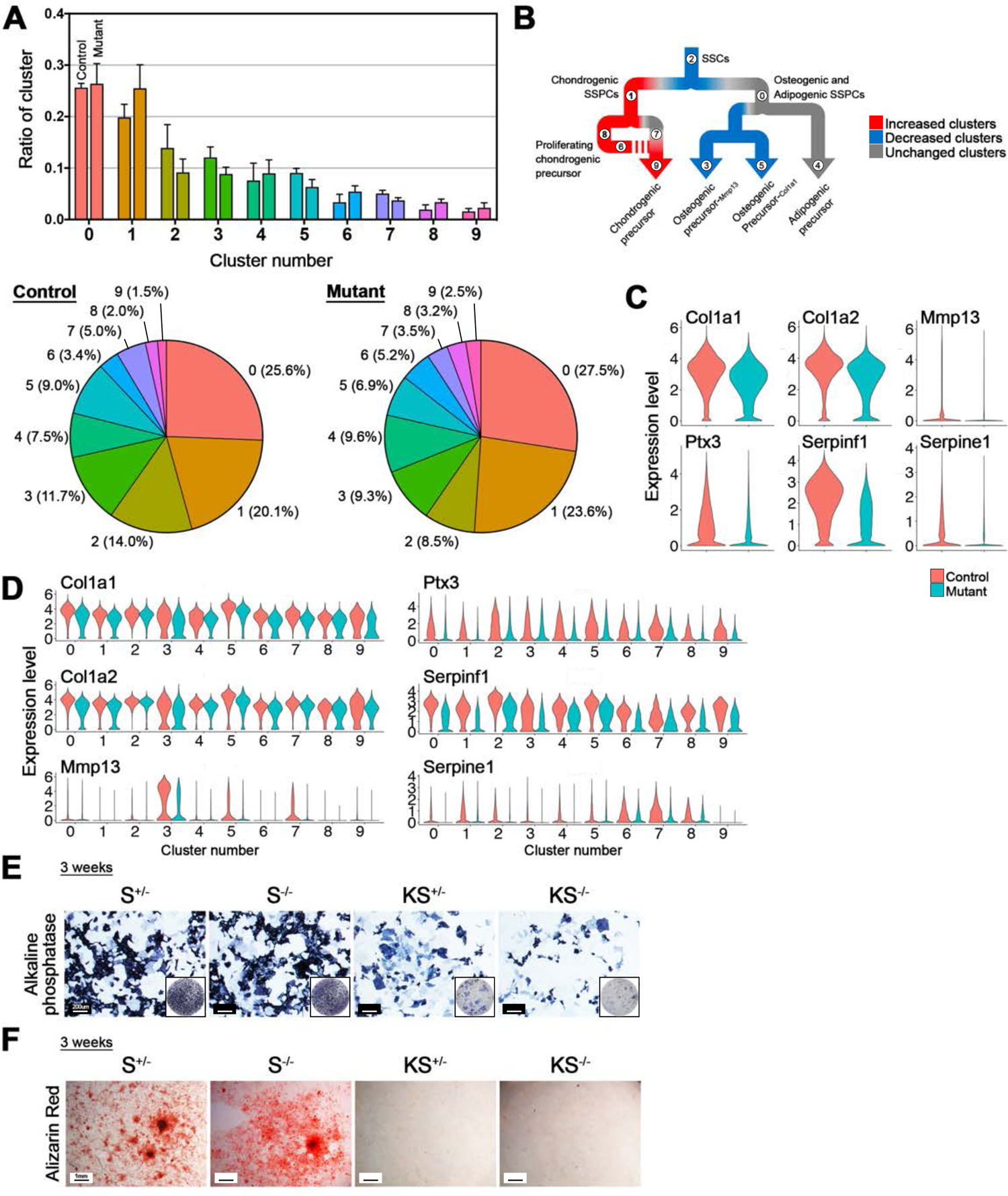
PDGFR*β*^D849V^ impairs osteogenesis in SSCs and SSPCs *in vitro*. **(A)** Integrated scRNA data with all four genotypes after subgrouping into two categories, control (*S^+/-^* and *S^-/-^*) and mutant (*KS^+/-^* and *KS^-/-^*) in Seurat. The proportional ratio of each cluster in controls and PDGFR*β* mutants were plotted (above) and also presented as pie charts with ratio labelling (below). **(B)** Color-coded PDGFR*β* mutant skeletal lineages with their increase (red), decrease (blue) and no change status (gray) based on Figure 4A’s ratio. **(C)** Expression of the top 6 significantly downregulated osteogenesis-related genes (*Col1a1*, *Col1a2*, *Mmp13*, *Ptx3*, *Serpinf1* and *Serpine1*) in mutant colonies. **(D)** Expression of the top 6 genes shown in (C) throughout 10 clusters between controls and mutants. **(E)** Primary SSPCs stained for alkaline phosphatase after 2 weeks of osteoblast differentiation (n = 3 biological replicates per genotype). **(F)** SSPCs stained with alizarin red after 4 weeks of osteoblast differentiation (n = 3 biological replicates per genotype).

To examine PDGFR*β*-mediated osteogenic defects *in vitro*, we isolated primary SSPCs from 3-week-old long bones (see Methods) and cultured them for osteoblast differentiation with a standard cocktail of inducers. *KS^+/-^* and *KS^-/-^* SSPCs showed reduced alkaline phosphatase staining, which indicates defective osteoblast differentiation (Figure 4E), and reduced alizarin red staining, which indicates defective mineralization (Figure 4F).

Next, to examine bone mass and mineralization in mice, we used micro-computed tomography (microCT) to examine 3-week-old tibias from the four original genotypes. *KS^+/-^* bones displayed osteoporosis-like pores in the cortical bone of the diaphysis (Figure 5A) and reduced trabecular bone formation in the proximal tibial metaphysis (Figure 5B). *KS^-/-^* bones had no pores and partially normalized bone mass and mineralization in both cortical and trabecular regions (Figure 5A and 5B). However, 5 to 8-week-old KS^-/-^ bones displayed less bone mass and mineralization in the cortical bone of the diaphysis (Figure 5C) and the proximal tibial metaphysis (Figure 5D) compared to age-matched control bones. To corroborate PDGFR*β*^D849V^-mediated bone formation defects, we performed calcein double staining to quantify mineral appositional growth rate (MAR). Histomorphometric analysis of cortical bone at 3 weeks showed decreased bone growth in *KS^+/-^*, which was normalized in *KS^-/-^* (Figure 5E). Interestingly, 6-week-old *KS^-/-^* cortical bone showed increased MAR compared to controls (Figure 5F), but the calcein labelling was thicker and more diffuse than controls, which is consistent with incomplete or delayed mineralization. In summary, as suggested by transcriptomics, we find that PDGFR*β*^D849V^ impairs osteogenic differentiation in cells derived from *KS^+/-^* and *KS^-/-^* mice, and leads to osteopenia that becomes severe early in *KS^+/-^* mice (by 3 weeks) and later in *KS^-/-^* mice (by 6 weeks).

**Figure 5.**
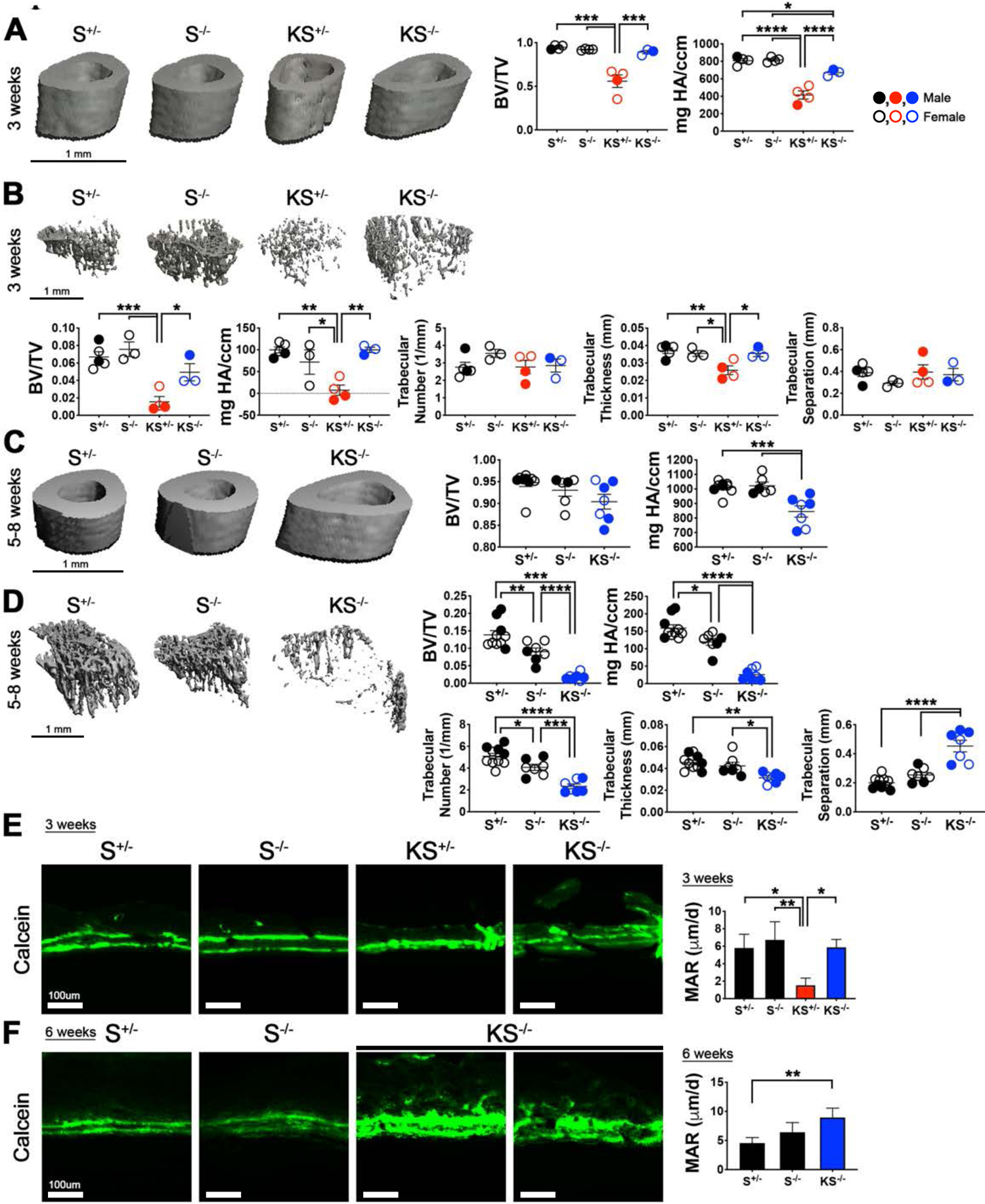
PDGFR*β*^D849V^ inhibits mineralization *in vivo*. **(A)** 3-week-old cortical bone mass and mineral density measured by microCT (n = 4 for *S^+/-^*, *S^-/-^* and *KS^+/-^*, n = 3 for *KS^-/-^*). **(B)** 3-week-old trabecular bone mass and mineral density (n = 5 for *S^+/-^*, 3 for *S^-/-^*, 4 for *KS^+/-^* and 3 for *KS^-/-^*). **(C)** 5-8-week-old cortical bone mass and mineral density (n = 9 for *S^+/-^*, 6 for *S^-/-^* and 7 for *KS^-/-^*). **(D)** 5-8-week-old trabecular bone mass and mineral density (n = 10 for *S^+/-^* and 7 for *S^-/-^* and *KS^-/-^*). **(E-F)** Histomorphometric analysis of calcein double-labelled proximal periosteal femur of four genotypes and mineral apposition rate (MAR) quantified. Calcein was injected twice into 3-week-old (D) or 6-week-old mice (E) with a 4-day interval. Mice were harvested 2 days after the second calcein injection (n = 3 per genotype for 3 weeks, n = 3-5 for 6 weeks). ****, *p* < 0.001, ***, *p* < 0.005, **, *p* < 0.01 and *, *p* < 0.05 by one-way ANOVA. Data represent mean ± SEM.

### PDGFRβ^D849V^ and STAT1 augment chondrogenic fate of SSCs

The growth and survival phenotypes (Figure 1A), and the time required to develop osteopenic phenotypes (Figure 5) are very different between *KS^+/-^* and *KS^-/-^* mice. This indicates a strong modifier effect of *Stat1*, at least in part due to *Stat1*-mediated autoinflammation downstream of PDGFR*β*^D849V^ (He et al. 2017). To identify changes in cell clusters and gene expression contributed by *Stat1*, we split the scRNA data into four groups representing the original genotypes *S^+/-^*, *S^-/-^*, *KS^+/-^* and *KS^-/-^* (Supplementary Fig. 5A and 5B, Supplementary Table 6). We found that *KS^+/-^* decreased SSCs (cluster 2), but prominently increased chondrogenic SSPCs (cluster 1), proliferating chondrocytes (cluster 6 and 8) and *Acan^high^*-chondrogenic precursors (cluster 9) compared to two controls (Figure 6A, Supplementary Figure 5B and 5C). In comparison, *KS^-/-^* moderately increased the chondrogenic SSPCs (cluster 1) and proliferating chondrogenic precursors (cluster 8), but less than *KS^+/-^*. We further analyzed differentially expressed genes specific to the *KS^+/-^* genotype (Supplementary Figure 5B and 5D). Among 34 genes specifically upregulated in *KS^+/-^* versus the other three genotypes, cartilage development genes were highly enriched, including *Sox9*, *Col2a1*, *H19* and *Acan* (Figure 6B, Supplementary Table 3 and 4). Increased *Sox9*, *Col2a1* and *H19* expression was not limited to cluster 9, but also showed increased expression in other clusters (Figure 6C, Supplementary Figure 5E). *Col2a1* and *H19* are known downstream targets of Sox9, which is a master transcription factor for chondrocyte proliferation and differentiation (Akiyama et al. 2002) and directly regulates collagen 2 production (Bell et al. 1997). Sox9 also indirectly promotes collagen 2 expression via a long non-coding RNA *H19* and its micro-RNA, *miR675* (Dudek et al. 2010). This suggests that increased Sox9 in *KS^+/-^*SSCs promotes chondrogenic proliferation and commitment. To evaluate chondrogenesis *in vivo*, we analyzed Safranin-O-stained tibias at 3 weeks old. Although *KS^+/-^* tibias were smaller in size, the proportional area of *KS^+/-^* cartilage was larger than the three other genotypes (Figure 6D). *KS^-/-^* also mildly increased cartilage area. We evaluated Sox9 expression *in vivo* in femurs from 3-week-old mice. Sox9-positive cell numbers were increased in both *KS^+/-^* and *KS^-/-^* genotypes, each with a distinct distribution in the distal femur. *KS^+/-^* showed ectopic expansion of Sox9-positive cells near the growth plate (Figure 6E). *KS^-/-^* also increased Sox9-positive chondrocytes in the proliferating and prehypertrophic zones within the growth plate (Figure 6E), consistent with the expansion of proliferating chondrogenic precursors (clusters 1 and 8) (Supplementary Figure 5D). To evaluate *KS^+/-^* chondrogenesis in autoinflammation-free conditions, we performed pellet culture with primary SSPCs from 3-week-old long bones of the four genotypes. We found that *KS^+/-^* SSPCs dramatically increased pellet size compared to the others after 21 days in culture (Figure 6F). Interestingly, while many *KS^+/-^* cells appeared to have matured into chondrocytes in matrices that were strongly positive for toluidine blue and safranin-O, cells in the pellet core appeared to remain undifferentiated (Figure 6F). *KS^+/-^* cells often generated irregular donut shaped pellets, unlike spherical pellets formed by control cells. *KS^-/-^* cells generated chondrocyte pellets similar to controls. Together, these findings suggest that PDGFR*β*^D849V^ and STAT1 enhance chondrogenic fate within the SSC lineage.

**Figure 6.**
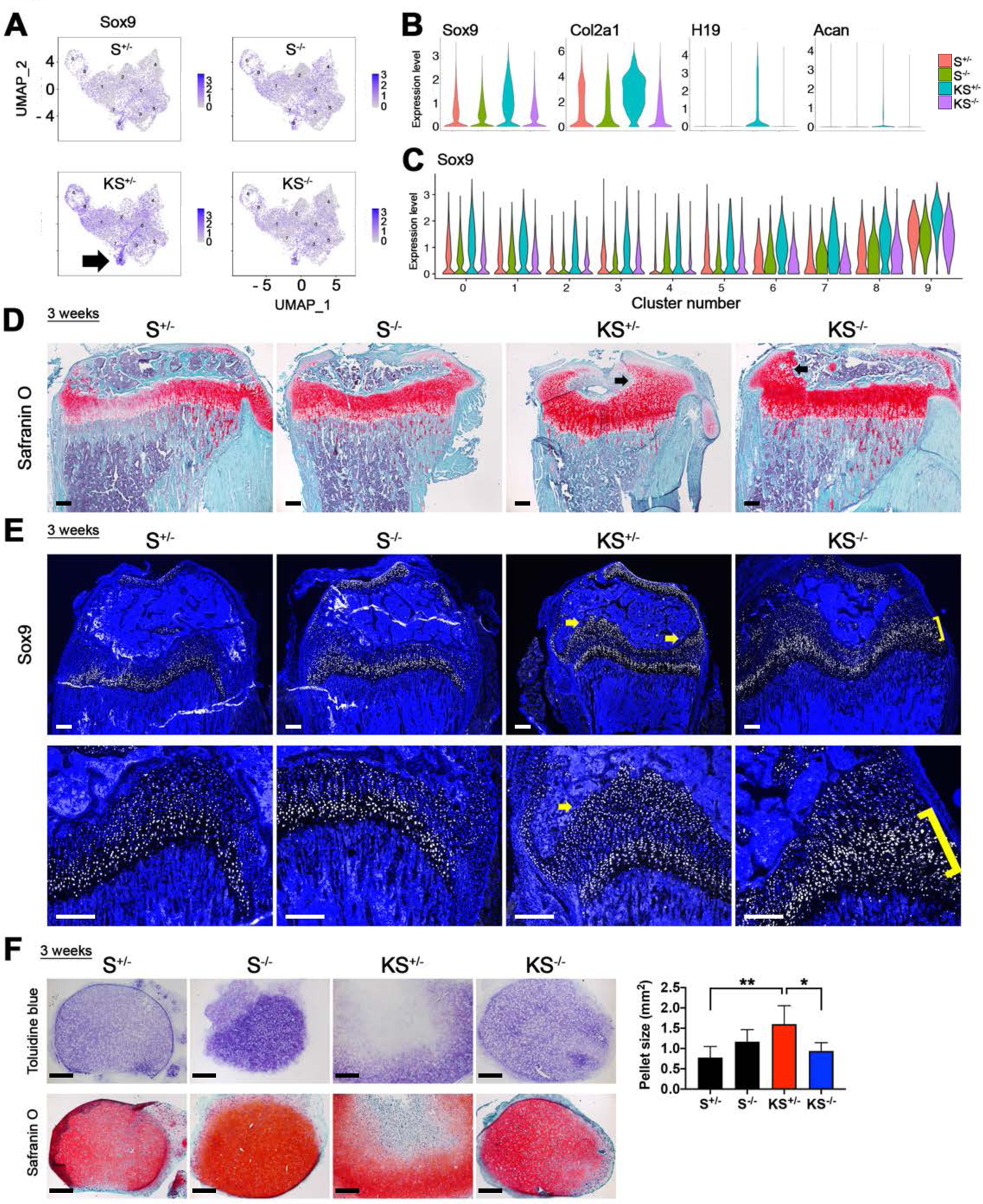
PDGFR*β*^D849V^ augments chondrogenic fate. **(A)** Distribution of Sox9 expressing cells in UMAP between genotypes. Arrow indicates *Acan^high^* chondrogenic precursors (cluster 9) **(B)** Expression levels of chondrogenic markers *Sox9*, *Col2a1*, *H19* and *Acan* in scRNA colonies between four genotypes. **(C)** Expression levels of *Sox9* throughout 10 clusters between four genotypes. **(D)** Safranin-O stained 3-week-old tibias of four genotypes with fast green counterstain (n = 3 for *S^+/-^*, *S^-/-^* and *KS^+/-^* and 6 for *KS^-/-^*). Black arrows indicate cartilage expansion. **(E)** Immunostaining of SOX9 in 3-week-old femurs (representative of n = 3 for each genotype). Ectopic SOX9 distribution indicated by yellow arrows in *KS^+/-^* and yellow brackets in *KS^-/-^*. **(F)** Histological analysis of Toluidine blue-stained or Safranin-O-stained chondrocyte pellets generated from SSPCs after 3 weeks of differentiation. The size of pellets was quantified with the histological sections of four genotypes (n = 5 biological replicates per genotype). Scale bar, 200 μm. **, *p* < 0.01 and *, *p* < 0.05 by one-way ANOVA. Data represent mean ± SEM.

### Low interferon signature in SSCs

Since *KS^+/-^* mice showed wasting due to PDGFR*β*-STAT1-mediated auto-inflammation, and *KS^+/-^* intrinsically overexpress interferon-stimulated genes (ISGs) in dermal fibroblasts (He et al. 2017) and SSPCs (Supplementary Figure 6A), we examined whether *KS^+/-^* SSCs also exhibit an ISG signature. A few ISGs, including *Ifi27*, *Bst2*, *Isg15*, *Ifit1*, *Psmb8*, *Mx1,* and *Stat1*, were upregulated in *KS^+/-^* SSCs throughout all clusters (Supplementary Figure 6B-6D). However, expression was very low for most ISGs and was detected in few cells of the SSC colonies regardless of genotype. It has been shown that other types of stem cells are intrinsically protected from interferon responses (Burke et al. 1978; Eggenberger et al. 2019), and this may be true as well for SSCs.

### PDGFRβ^D849V^ activates STAT5 and increases *Igf1* expression

Although both *KS^+/-^* and *KS^-/-^* mutants showed defective mineralization, only *KS^-/-^* showed overgrowth *in vivo*. From scRNA data, we found that *Igf1* and *Socs2* were highly upregulated in *KS^-/-^* SSC colonies and were upregulated to a lesser amount in *KS^+/-^* colonies, compared to controls (Figure 7A). Increased *Igf1* and *Socs2* mRNAs were detected in most *KS^+/-^* and *KS^-/-^* clusters, from SSCs to precursors (Figure 7B). *Igf1* and *Socs2* are direct transcriptional targets of STAT5. The STAT5-IGF1 axis is critical for the biological effects of growth hormone receptor signaling (Udy et al. 1997; Chia et al. 2006), and SOCS2 is involved in negative feedback on STAT5 (Greenhalgh et al. 2002). It is known that PDGFR*β* can phosphorylate STAT5 (Valgeirsdottir et al. 1998), although the biological significance is unclear. Interestingly, *Stat5a* mRNA was overexpressed by *KS^+/-^* colonies and *Stat5b* was overexpressed by *KS^-/-^* colonies (Figure 7C). However, the role of STAT5 in signaling and gene activation is mainly regulated by phosphorylation. Both *KS^+/-^* and *KS^-/-^* SSC colonies exhibited constitutive STAT5 phosphorylation (Figure 7D), and STAT5 was constitutively phosphorylated in SSPCs from both mutants (Figure 7E). The antibodies used for these Western blots detect both STAT5 proteins and therefore cannot discern whether there is differential activation of the two isoforms in *KS^+/-^* and *KS^-/-^* colonies. Based on these results, we suggest that STAT5-IGF1 axis could be a key signaling pathway for PDGFR*β*-mediated overgrowth. Serum IGF1 levels were very low in 3-week-old *KS^+/-^* mice, which were moribund. Serum IGF1 was unchanged between controls and *KS^-/-^* mice, which were on the cusp of overgrowth (Figure 7F). However, IGF1 levels were elevated in culture medium of *KS^+/-^* and *KS^-/-^* SSPCs (Figure 7G). Therefore, PDGFR*β* mutant skeletal lineages may promote skeletal overgrowth by increasing IGF1 levels locally, rather than systemically. We suggest that both *KS^+/-^* and *KS^-/-^* mice have potential for overgrowth due to increased signaling through the STAT5-IGF1 axis. But *KS^+/-^* mice do not exhibit the overgrow phenotype because of growth-suppressive (and lethal) effects of *Stat1*-mediated autoinflammation.

**Figure 7.**
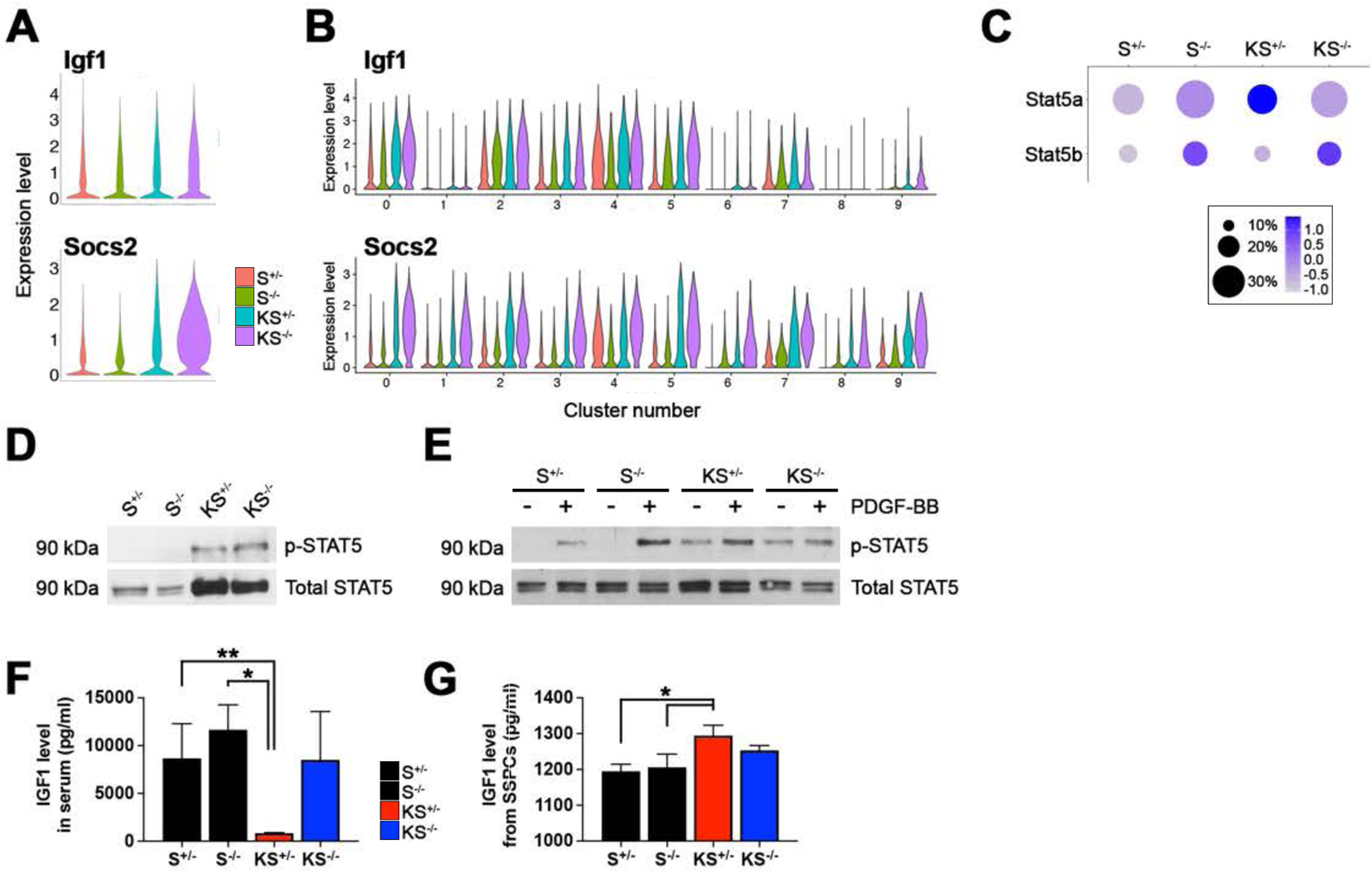
PDGFR*β*^D849V^ enhances growth signaling through STAT5 and IGF1. **(A)** Expression levels of *Igf1* and *Socs2* in scRNA colonies between four genotypes. **(B)** Expression levels of *Igf1* and *Socs2* throughout 10 clusters between four genotypes. **(C)** Expression levels and percent expression of *Stat5a* and *Stat5b* between four genotypes in all clusters of scRNA colonies. **(D)** Phosphorylated STAT5 (pY694) and total STAT5 in SSC colonies of four genotypes after overnight serum-starvation, representative of two biological replicates. **(E)** Phosphorylated STAT5 (pY694) and total STAT5 in primary SSPCs of four genotypes after overnight serum-starvation PDGF-BB, representative of three biological replicates. **(F)** IGF1 levels in serum collected from 3-week-old mice of four genotypes (n = 5 for *S^+/-^*, *S^-/-^* and *KS^-/-^* and 3 for *KS^+/-^*). **(G)** IGF1 levels in cell culture supernatants collected from SSPCs (n = 3 for *S^+/-^*, *S^-/-^* and *KS^-/-^* and 4 for *KS^+/-^*). **, *p* < 0.01 and *, *p* < 0.05 by one-way ANOVA. Data represent mean ± SEM.

## Discussion

The depletion of SSCs by conditional diphtheria toxin expression in mice shows the importance of SSCs in skeleton growth (Worthley et al. 2015; Mizuhashi et al. 2018), but signaling pathways and genes regulating SSC functions are still largely unknown. In this study, we found that increased PDGFR*β* signaling alters SSC abundance and lineage differentiation in parallel to phenotypes affecting the skeleton. Our results suggest that constitutive PDGFR*β* signaling in SSCs precedes alterations in osteogenic and chondrogenic differentiation. These cell-autonomous and skeletal lineage-autonomous defects, combined with systemic effects of constitutive PDGFR*β* signaling (including interferonopathy-like autoinflammation), lead to osteopenia and skeletal phenotypes that are reminiscent of humans with *PDGFRB* mutations.

Our work demonstrates the utility of using scRNA sequencing to uncover pathways and genes in cultured stem cells to provide new molecular understanding of diseases. Colony formation assays have long been used to assay for the enrichment of cells with stem cell properties, but it has been challenging to identify molecular targets or cellular pathways due to the rarity of SSCs. In our case, it was important to isolate SSCs and culture them because the *KS^+/-^* mutants exhibit severe autoinflammation *in vivo.* With scRNA transcriptomics of cultured SSCs, we overcame these limitations and identified sub-populations of SSC and precursor cells with alterations that are linked to phenotypes in *KS^+/-^* and *KS^-/-^* mutant mice.

We found that wasting *KS^+/-^* mutants and overgrown *KS^-/-^* mutants both exhibit impaired osteogenesis *in vivo*. Both mutant SSC colonies decreased osteogenesis-related genes such as *Col1a1, Col1a2,* and *Serpinf1* and both mutant SSPCs showed reduced osteogenic differentiation and mineralization capacities. Similar to mice, humans show osteogenesis imperfecta caused by loss of function mutations in *COL1A1* or *COL1A2* (MIM 166200) or *SERPINF1* (MIM 172860) with decreased mineralization and brittle bones (Byers and Pyott 2012). However, differentially expressed genes between *KS^+/-^* and *KS^-/-^* mutants suggest that *Stat1*-depedent mechanisms are also involved. We highlighted increased *Sox9* expression in *KS^+/-^* mice and SSC colonies as an intrinsic source of osteogenesis defects because *Sox9* overexpression in mice leads to impaired osteogenesis and dwarfism (Akiyama et al. 2004; Zhou et al. 2006). *KS^+/-^* osteogenesis defects *in vivo* are likely compounded by autoinflammation (Redlich and Smolen 2012). The kidneys are critical for maintaining healthy bones by regulating phosphorous and calcium levels in the blood. A PDGFR*β* gain-of function mutation controlled by *Foxd1Cre^tg^* has been shown to cause mesangioproliferative glomerulonephritis and renal fibrosis with progressive anemia in mice (Buhl et al. 2020). However, as these phenotypes begin to appear at 6 weeks of age, renal dysfunction is unlikely to initiate skeletal phenotypes in our mice, but it may contribute to later progression in *_KS_-/-*.

Our study suggests that *Pdgfrb^D849V^* signaling through STAT1 promotes *Sox9* expression and chondrocyte proliferation. We found that *KS^+/-^* SSPCs formed irregularly shaped chondrocyte pellets. The non-spherical morphology may occur because of excessive cartilage matrix. For example, bone marrow stromal cells obtained from familial osteochondritis dissecans (FOCD) patients (*ACAN* loss-of-function mutant) produce enlarged and irregularly shaped chondrocyte pellets with highly upregulated cartilage matrix proteins including collagen 11 and proteoglycans (Xu et al. 2016). We found that *Acan*, *Col2a1,* and *Col11a1* were all upregulated in *KS^+/-^* SSC colonies. We do not know how *Pdgfrb^D849V^*-STAT1 regulates *Sox9*. But STAT3, a related transcription factor, has been shown to directly regulate *Sox9* in chondrogenesis (Hall et al. 2017), and STAT1 can heterodimerize with STAT3.

SSCs with *Pdgfrb^D849V^* increase functional SSC numbers and growth signaling gene expression (ie., *Igf1* and *Socs2*). STAT5 may be an important mediator of overgrowth because *Igf1* is a direct transcriptional target of STAT5, which is directly activated by PDGFR*β* (Valgeirsdottir et al. 1998; Chia et al. 2006). STAT5 activation is constitutive in mutant SSCs and SSPCs, but only *KS^-/-^* induces overgrowth. As discussed above, there are additional effectors in *KS^+/-^* to counterbalance potential overgrowth, such as STAT1-dependent autoinflammation and *Sox9* expression. Furthermore, *KS^+/-^* SSCs upregulated the expression of IGF binding proteins (IGFBP3, 6 and 7) (Supplementary Table 3 and 4), potentially blocking IGF1 signals. Increased IGF1 in *KS^-/-^* mice seems to occur locally in SSCs and SSPCs rather than systemically because increased IGF1 was detected in mutant SSPC conditioned medium but not in mutant mouse circulation. We do not know if IGF1 causes overgrowth through autocrine signaling or by signaling between cell types. Tissue-specific genetic studies will be needed to investigate the involvement of a PDGFR*β*-STAT5-IGF1 axis and to identify IGF1-responsive cells.

Although the D849V mutation in our study is not the same as mutations seen in Penttinen syndrome and Kosaki overgrowth syndrome, we believe our findings here are closely related to the human conditions. Osteopenia, often associated with bone fractures, is a common feature in both human diseases. For example, two Kosaki overgrowth syndrome patients developed fractured tibias and compression fractures in the spine, causing deformation of their bones (Minatogawa et al. 2017; Foster et al. 2020). Further, a Penttinen Syndrome patient exhibited osteoporosis with multiple fractures (Johnston et al. 2015). Similarly, we found osteopenia in *Pdgfrb^D849V^* mice regardless of their phenotype on the wasting-overgrowth spectrum. A better understanding of the pathogenic mechanisms of different activating PDGFR*β* mutations will be obtained through future genetic models that reproduce the specific human *PDGFRB* mutations seen in Penttinen and Kosaki syndromes. The current work puts forth SSCs as a conceptual tool for considering pathological changes in the PDGFR*β* mutant skeleton as resulting from altered stem cell functions.

## Materials and Methods

### Animal models

Mouse strains *Pdgfrb^D849V^* (#018435), *Sox2Cre^tg^* (#008454) and *Stat1^flox^* (#012901) are available at the Jackson Laboratory. All procedures performed on mice were approved by the Institutional Animal Care and Use Committee of the Oklahoma Medical Research Foundation. Mice were maintained on a mixed C57BL/6;129 genetic background with a standard mouse chow diet (5053 Purina). Mutant *KS^+/-^* and *KS^-/-^* mice were compared with age and sex-matched littermate control *S^+/-^* and *S^-/-^* mice.

### Fluorescence-activated cell sorting (FACS) and CFU-F of skeletal stem cells

PDGFR*α* and Sca1 double positive SSCs (P*α*S cells) were isolated by the method of Houlihan et al. (Houlihan et al. 2012) with some modifications. FACS buffer was made of 1x HBSS (Gibco), 2% FBS, 1x penicillin/streptomycin (Gibco), 1 mM ethylenediaminetetraacetic acid (EDTA, VWR) in autoclaved H_2_O. Limbs of control and mutant mice at 3, 6 and 9 weeks old were dissected and soaked in 70% EtOH for 2 mins, followed by careful removal of adherent muscles. Bone marrow (BM) was flushed 2-3 times with sterile PBS using a 26G x 1/2 needle and 1 ml syringe (BD). BM-free bones were cut into small pieces with sterile scissors until bones became paste (usually 3 mins for 3 weeks old and 5 mins for 6-9 weeks old). The bone paste from each mouse was transferred to a 15 ml conical tube and enzymatically digested with 15 ml 0.2% type II collagenase (Worthington) in DMEM (Corning) for 1 hour with agitation at 37 °C. Digested bone paste was filtered through 70 μm cell strainer and collected in a 50 ml conical tube on ice. Bone paste remaining on the filter was collected with 2.5 ml FACS buffer, gently tapped in a mortar with a pestle and filtered into the same 50 ml conical tube on ice. This was repeated until the total volume of filtrate reached 50 ml. After centrifugation at 280 x *g* for 10 min at 4 °C, 1 ml ice-cold sterile H_2_O was used to remove red blood cells for 6 sec. Then 1 ml 4% FBS in PBS and 13 ml FACS buffer were added, followed by filtering with 70 μm cell strainer into a new 15 ml tube. After centrifugation for 5 min, re-suspended cells were incubated with fluorophore-conjugated antibodies (Supplementary Table 8) on ice for 20 min in dark. P*α*S cells were sorted using MoFlo XDP Cell Sorter (Beckman Coulter) or FACSAria III (BD) by negative gating with Zombie Green^TM^(live/dead cell dye), CD31, CD45 and TER119 and positive gating with PDGFR*α* and Sca1 (Supplementary Figure 2). For colony formation assays, 2,000 P*α*S cells were cultured in a 6-well plate with 20% FBS (mesenchymal stem cell-qualified, Gibco), 25 units/mL penicillin/streptomycin and 2 mM L-glutamine in alpha MEM (Gibco) for 2 weeks under hypoxia (5% oxygen) at 37 °C. Colonies were fixed with 4% PFA for 5 min and stained with 0.5% crystal violet for quantification.

### Single-cell RNA sequencing and annotation

Single-cell suspensions of P*α*S colonies were harvested from CFU-F assays after culturing for 2 weeks as explained above. Trypsinized cells were filtered through a 40 μm cell strainer. 4 batches were generated in duplicate with 1 male and 1 female of each genotype. A total of 8 samples (maximum 20,000 cells per each sample) were loaded for the 10X Genomics Chromium platform, barcoded, and sequenced on the Illumina NovaSeq SP flowcell platform at a depth of 400 million reads per sample. See Supplementary Table 5 for actual cell numbers sequenced and analyzed. Using the 10X Genomics Cellranger software (version 3), sequences were demultiplexed to extract cellular barcodes, cDNA inserts and unique molecular identifiers (UMIs) and the cDNA sequences were aligned to the murine mm10 genome reference to count UMIs. Quality control, data integration, clustering and gene expression analysis were performed with Seurat package V3 for R (Butler et al. 2018; Stuart et al. 2019). Cells with > 8,000 genes, < 1,500 genes, or > 8% of genes mapping to mitochondrial genes were removed as poor quality or doublets. Eight samples were individually normalized by using the log transformation method in Seurat. Data integration of 8 samples to remove batch effect was done with canonical component analysis (CCA) with the top 2,000 variable genes identified by FindVariableFeatures function in Seurat (Butler et al. 2018). Cells were categorized into S, G1 or G2/M phase by scoring cell cycle-associated gene expression (Kowalczyk et al. 2015). Principal component analysis (PCA) was performed by regressing out cell cycle, mitochondrial and ribosomal gene expression. The top 100 PCs were selected to perform dimensional reduction using uniform manifold approximation and projection (UMAP). We determined a cluster resolution of 0.4 (total 12 Clusters) using Clustree R package (graphic-based cluster resolution analyzer) (Zappia and Oshlack 2018). Among the 12 clusters, 10 were selected for analysis, because 2 of the clusters only included about 10 cells per each. A heatmap was generated with the top 10 genes (Supplementary Table 1) in each cluster determined by FindAllMarkers function in Seurat. For gene signature analysis, differentially expressed genes (DEGs) of each cluster were determined by FindMarkers function in Seurat with 0.25 log-scale fold-change threshold between two groups (ie., genotypes). DEGs were further analyzed with the Database for Annotation, Visualization and Integrated Discovery (DAVID) to determine gene ontology and pathway identification (Huang da et al. 2009). Cell types of clusters were defined with a signature gene list, gene ontology and literature as SSCs, SSPCs, chondrocyte precursors, osteoblast precursors and adipocyte precursors. Trajectory analysis was performed using Slingshot (Street et al. 2018). SingleCellExperiment objects (transformed from Seurat objects) and UMAP information were utilized as input for trajectory predictions. We selected cluster 2 (SSCs) as the top of the hierarchy. Sub-grouping and trajectory analysis with chondrogenic lineages (clusters 1, 2, 6, 7, 8 and 9) were additionally performed to determine an additional branch (clusters 8>6>9). To identify PDGFR*β* GOF-specific genes and clusters, we split the integrated data into 2 groups (controls *S^+/-^* and *S^-/-^* versus mutants *KS^+/-^* and *KS^-/-^*) or 4 groups (individually *S^+/-^*, *S^-/-^*, *KS^+/-^* and *KS^-/-^*). DEGs between 2 or 4 groups were identified with by FindMarkers function in Seurat.

### Primary skeletal stem and progenitor cell (SSPC) culture and tri-lineage differentiation

Compact bone was used to isolate primary skeletal stem and progenitor cells (Zhu et al. 2010). BM-free bones were collected as described above and were cut into small pieces (1-3 mm^3^) with sterile scissors. The bone chips were transferred into a 1.6 ml microcentrifuge tube and enzymatically digested with 1.5 ml of 1 mg/ml type II collagenase (Worthington) in alpha MEM (Corning) plus 10% FBS (mesenchymal stem cell-qualified, Gibco) for 1.5 hours with agitation in a 37 °C incubator. The bone chips were washed with 1 ml of alpha MEM, seeded into a 6-well plate, and maintained with alpha MEM plus 10% FBS (mesenchymal stem cell-qualified, Gibco), 25 units/mL penicillin/streptomycin and 2 mM L-glutamine at 37 °C. 0.25 %. Passages 4 to 6 were used for tri-lineage differentiation assays. For osteogenesis, 2 x 10^5^ cells were seeded in a 24-well plate. On the following day, they were treated with alpha MEM supplemented with 10% FBS (mesenchymal stem cell-qualified), 10^-7^ M dexamethasone, 10 mM *β*-glycerol-phosphate and 50 mM ascorbate-2-phosphate. Medium was changed 3 times per week for 2 weeks (early osteoblast differentiation) and 4 weeks (mineralization). After 2 weeks, some samples were subjected to alkaline phosphatase (ALP) staining with ALP buffer (100mM Tris-HCl, 100mM NaCl, 5mM MgCl_2_, 0.05% Tween-20 in deionized water (adjusted at pH 9.5)) plus 0.02% 5-bromo-4-chloro-3-indolyl phosphate (BCIP) and 0.03% nitro blue tetrazolium (NBT). After 4 weeks, some samples were stained with 40 mM alizarin red. For chondrogenesis, 1 x 10^6^ cells were pelleted in a 15 ml conical tube at 280 x *g* for 6 min at 4 °C and cultured for 3 weeks with the MesenCult^TM^-ACF chondrogenic differentiation kit (Stemcell Tech). Pellets were fixed with 4% PFA for 1 hour, embedded in paraffin, sectioned at 8 μm and stained with 0.05% toluidine blue or 1% safranin-O. For SSC colony osteogenesis, colonies cultured for 2 weeks under hypoxia were further treated with the osteogenesis inducers as explained above for additional 2 weeks under normoxia and stained with alkaline phosphatase (Fast green as a counterstain). For SSC colony adipogenesis, 2-week-cultured colonies were treated with alpha MEM supplemented with 10% FBS (mesenchymal stem cell-qualified), 10^-6^ M dexamethasone, 250 μM IBMX, 10 μg/ml insulin and 5 nM rosiglitazone for 2 days. Colonies were maintained with alpha MEM plus 10% FBS (mesenchymal stem cell-qualified) and 10 ng/ml insulin for additional 8 days by changing 3 times in a week. Adipocytes were quantified with Oil-O-Red stain (Fast green as a counterstain).

### Enzyme-linked immunosorbent assay (ELISA)

IGF1 levels were measured in serum and SSPC culture supernatant using ELISA (Sigma-Aldrich, RAB-0229) according to the manufacturer’s protocol. Blood was collected from four genotypes at 3 weeks old, incubated at room temperature for 30 min, and centrifuged at 1,000 rpm for 10 min at 4 °C. Clear supernatant was stored at −20 °C until used. SSPCs (passages 4-5) isolated from four genotypes at 3 weeks old were cultured in a 24-well plate until confluent. After washing three times with sterile PBS, 500 μl serum-free fresh alpha MEM was replaced and cultured for 72 hrs. Cell culture supernatant was filtered through a 20 μm cell-strainer and stored at −20 °C until used. Both serum and cell culture supernatant were diluted to 1:100 for ELISA.

### Western blotting

For P*α*S SSC colonies, cells were starved in medium containing 0.2% FBS (mesenchymal stem cell-qualified) for 24 hours. For SSPCs, cells were starved in medium containing 0.1% FBS (mesenchymal stem cell-qualified) for 24 hours and treated with 10 ng/ml PDGF-BB (R&D systems) for 10-15 min. Cells were then lysed in RIPA buffer (50mM Tris pH 7.4, 1% NP-40, 0.25% sodium deoxycholate, 150 mM NaCl, 0.1% sodium dodecyl sulfate) with 1 mM NaF, Na_3_VO_4_, PMSF, and 1x protease inhibitor cocktail (Complete, Roche). Pierce BCA assay was utilized to determine protein concentration. Aliquots of 5-10 μg of protein were separated by SDS-PAGE, transferred to nitrocellulose membranes, blocked with 5% BSA, and incubated with primary antibodies (Supplementary Table 8) overnight at 4 °C. Membranes were probed with horseradish peroxidase (HRP)-conjugated secondary antibodies at 1:5000 (Jackson ImmunoResearch) in 5% milk. Pierce ECL Western blotting substrate (ThermoFisher) and autoradiography film (Santa Cruz) were utilized to develop blots.

### RNA isolation and qRT-PCR

Total RNA was isolated from cultured primary skeletal stem and progenitor cells enriched from compact bones using Trizol (ThermoFisher). cDNA reverse transcription was performed with Superscript III RT (Invitrogen) and random primers. Quantitative PCR was performed using a Bio-Rad iCycler with iQ TM SYBR green master mix and designated primers (Supplementary Table 7).

### Tissue and histology

Tissue was fixed overnight in 10% neutral buffered formalin (NBF, Sigma-Aldrich) or 4% paraformaldehyde (PFA, Electron Microscopy). Bones were decalcified in 0.5 M EDTA solution (pH 7.4) for 2 weeks at room temperature. For cartilage staining, fixed tissues were embedded in paraffin and sectioned at 6 μm thickness followed by Hematoxylin and Eosin (H&E) or Safranin O staining (Fast green and Hematoxylin as counter stain). For tartrate-resistant acid phosphate (TRAP) staining, fixed tissues were decalcified in 0.5 M EDTA solution at 4 °C, dehydrated in 20% sucrose in PBS and cryosectioned at 8-10 μm thickness followed by Naphthol AS-BI phosphoric acid and diazotized Fast Garnet GBC stains (Fast green as counter stain) (Sigma Aldrich, 387A). For immunostaining, PFA-fixed tissues were cryosectioned at 8-12 μm. Slides were stained with primary and secondary antibodies (Supplementary Table 8) and DAPI (Sigma) as a nuclear stain. Imaging was performed with a Nikon Eclipse 80i microscope with a digital camera.

### Bone formation rate and microcomputed tomography

To measure the mineral appositional growth rate (MAR) of bone formation, double calcein stain was performed. calcein (10 mg/kg, Sigma) in 0.9% NaCl and 2% NaHCO_3_ was intraperitoneally injected in 3- or 6-week-old mice with a 4-day interval. Mice were harvested 2 days after the second injection. Paraffin sectioning was performed as per the method of Porter *et al*. (Porter et al. 2017): femurs were harvested with adherent muscles and fixed with formalin for 2 days, then cleared with 10% KOH for 96 hours. The tissues were embedded in paraffin and sectioned at 6 μm thickness, pre-soaking the paraffin blocks in 1% KOH just before sectioning. For microCT, tibias were harvested and stored in 70% EtOH at 4 °C, then analyzed to measure bone volume (BV, mm^3^), total volume (TV, mm^3^), mean/density (mg HA/ccm), trabecular numbers (1/mm), trabecular thickness (mm), and trabecular separation (mm) with a VivaCT 40 microCT (Scanco Medical) at an X-ray tube volatage of 70 kV_p_ and an X-ray current 114 μA for 2- to 3-week-old bones or 55 kV_p_ and 85 μA for older bones.

### Image quantification and statistics analysis

The size of CFU-F and chondrocyte pellets, and the distance between double calcein stain, were quantified with ImageJ software (National Institute of Health). The Set Scale function was used to convert units from pixels to mm or μm with known scale of images. Regions of interest (ROIs) were selected using the polygon selection tool to measure the area of CFU-F and pellet images. The straight tool was utilized to measure the distance between double calcein stain. Statistical calculations were performed by using GraphPad Prism 9 with one-way or two-way ANOVA. Data were represented with mean ± standard error of the mean (SEM).

## Acknowledgements

We thank Jacquelyn C. Herron, Shouan Zhu, Mary Beth Humphrey, Timothy M. Griffin, and members of the Olson lab for their assistance and helpful discussion. We also thank the Microscopy Core and Mouse Phenotyping Core Facilities (associated with P30-GM114731) of the Oklahoma Medical Research Foundation Centers of Biomedical Research Excellence. H.R.K. was supported by F32-HL142222 from National Institutes of Health (NIH)/National Heart, Lung, and Blood Institute (NHLBI). This work was supported by US National Institutes of Health (NIH) grant R01-AR073828 (L.E.O.) and grants from the Oklahoma Center for Adult Stem Cell Research – a program of TSET (L.E.O).

## Author contributions

H.R.K designed experiments and analyzed data. H.R.K., J.P.W., and J.K. performed experiments. H.R.K. and L.E.O. wrote the manuscript. H.R.K., J.P.W., and L.E.O. edited the manuscript. All authors read and approved the final manuscript.

---

**Supplementary Figure 1.**
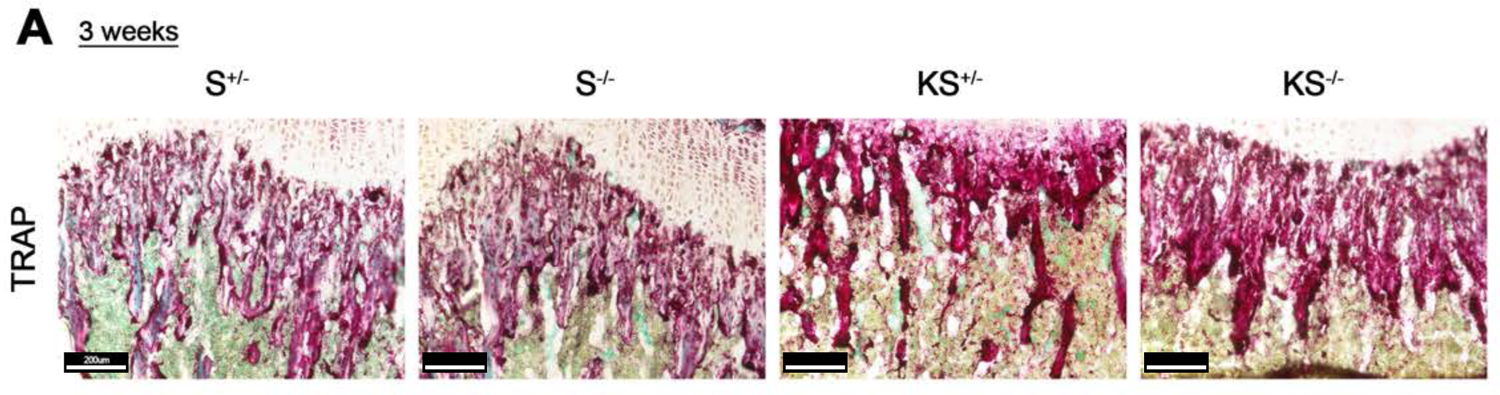
PDGFR*β* ^D849V^ increases osteoclasts. **(A)** TRAP-stained 3-week-old femurs of four genotypes. Fast green was used as a counterstain.

**Supplementary Figure 2.**
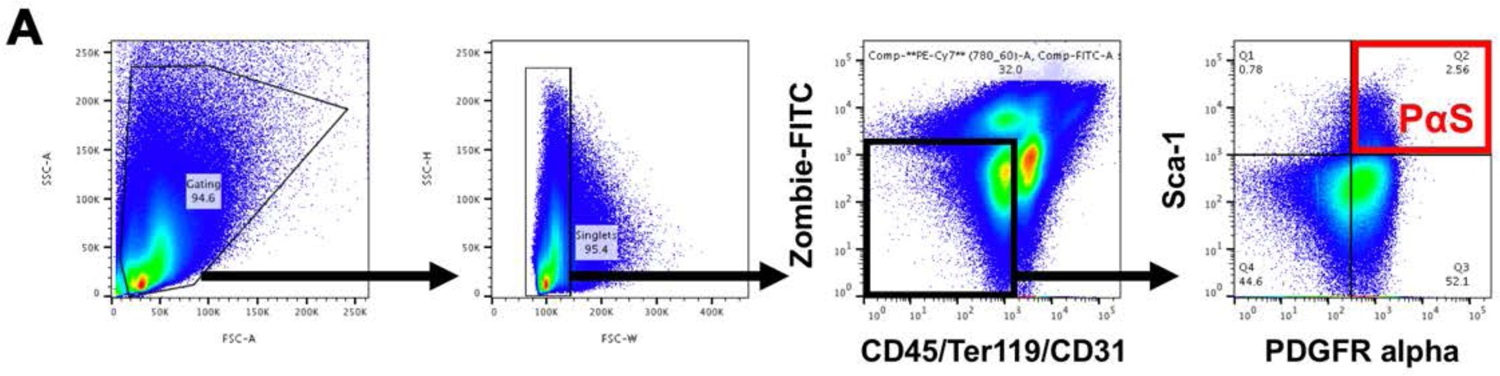
P*α*S SSCs sorting strategy. **(A)** After single-cell gating, SSCs were immunophenotyped or isolated by negative gating with Zombie-dye (live/dead cell dye), CD31, CD45 and TER119 and positive gating with PDGFR*α* and Sca1.

**Supplementary Figure 3.**
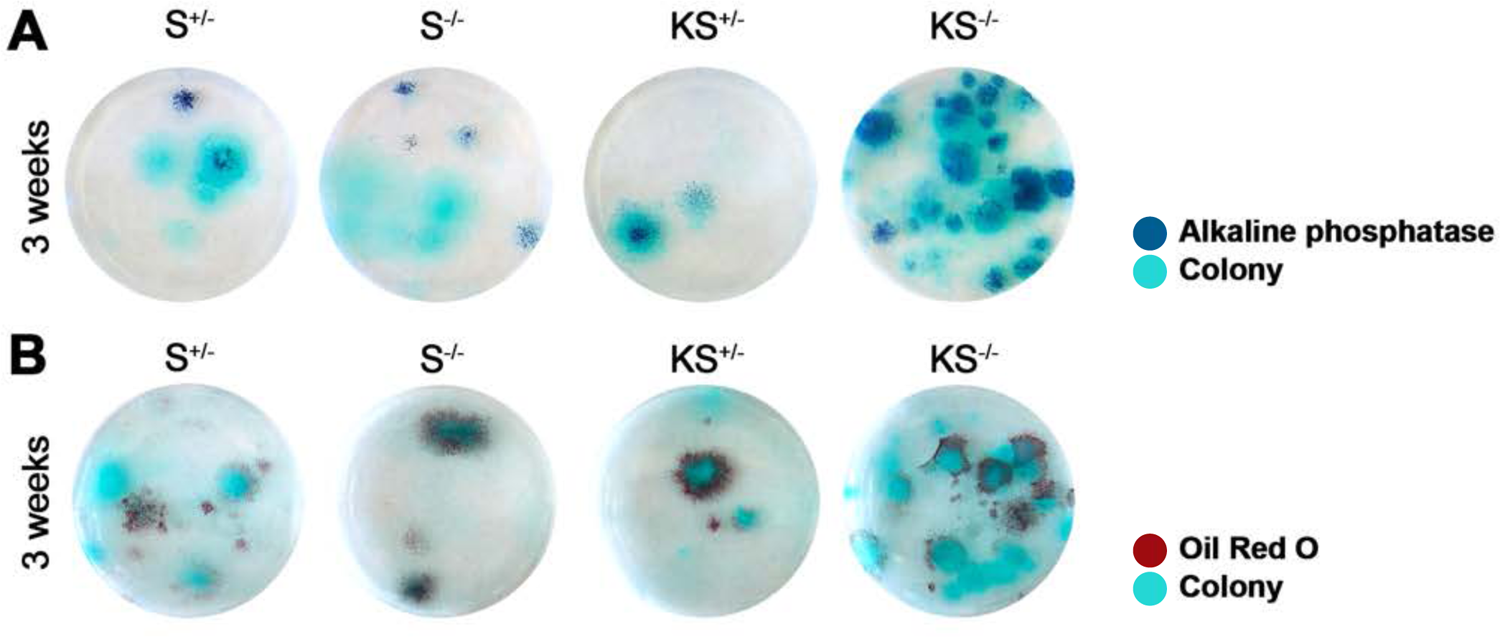
P*α*S SSC-derived colonies can differentiate into osteoblasts and adipocytes. **(A-B)** P*α*S SSCs were isolated from 3-week-old limbs and cultured for 2 weeks with 2,000 cells in a 6-well plate with hypoxia, followed by 2 weeks for osteogenesis (A) and adipogenesis (B). Alkaline phosphatase and Oil-Red-O were used to stain osteogenic and adipogenic cells, respectively. Fast green was used as a counterstain.

**Supplementary Figure 4.**
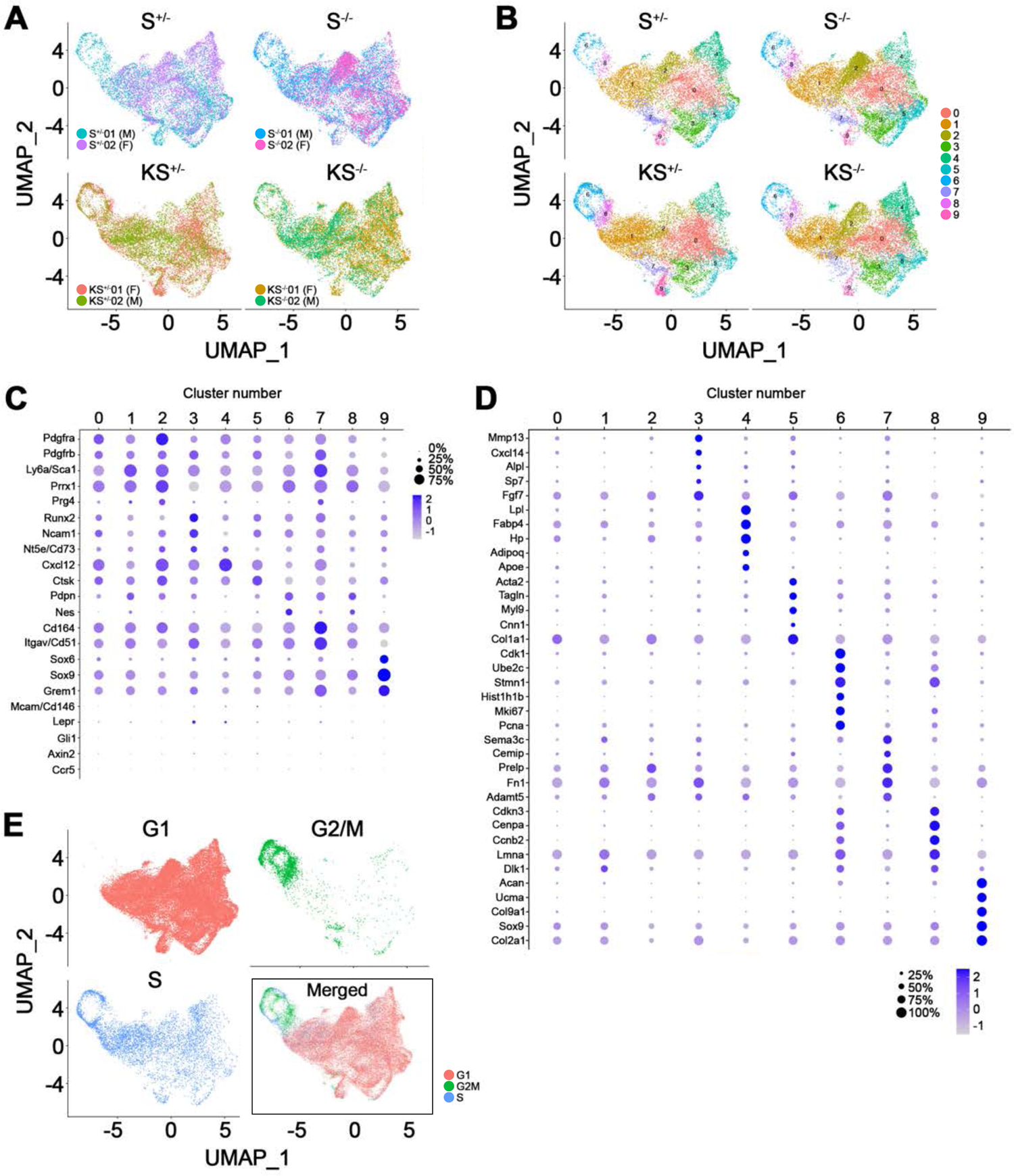
Single-cell RNA sequencing analysis of P*α*S SSC colonies. **(A)** UMAP plots of biological duplicates of each genotype. M and F indicates male and female mice, respectively. **(B)** UMAP plots of 10 cluster distributions for four genotypes. **(C)** Expression levels and percent expression of mesoderm, SSC and SSPC markers in each cluster. **(D)** Expression levels and percent expression of precursor markers from clusters 3 to 9. **(E)** UMAP plots of cell cycle phase (G1, G2/M, S and merged)

**Supplementary Figure 5.**
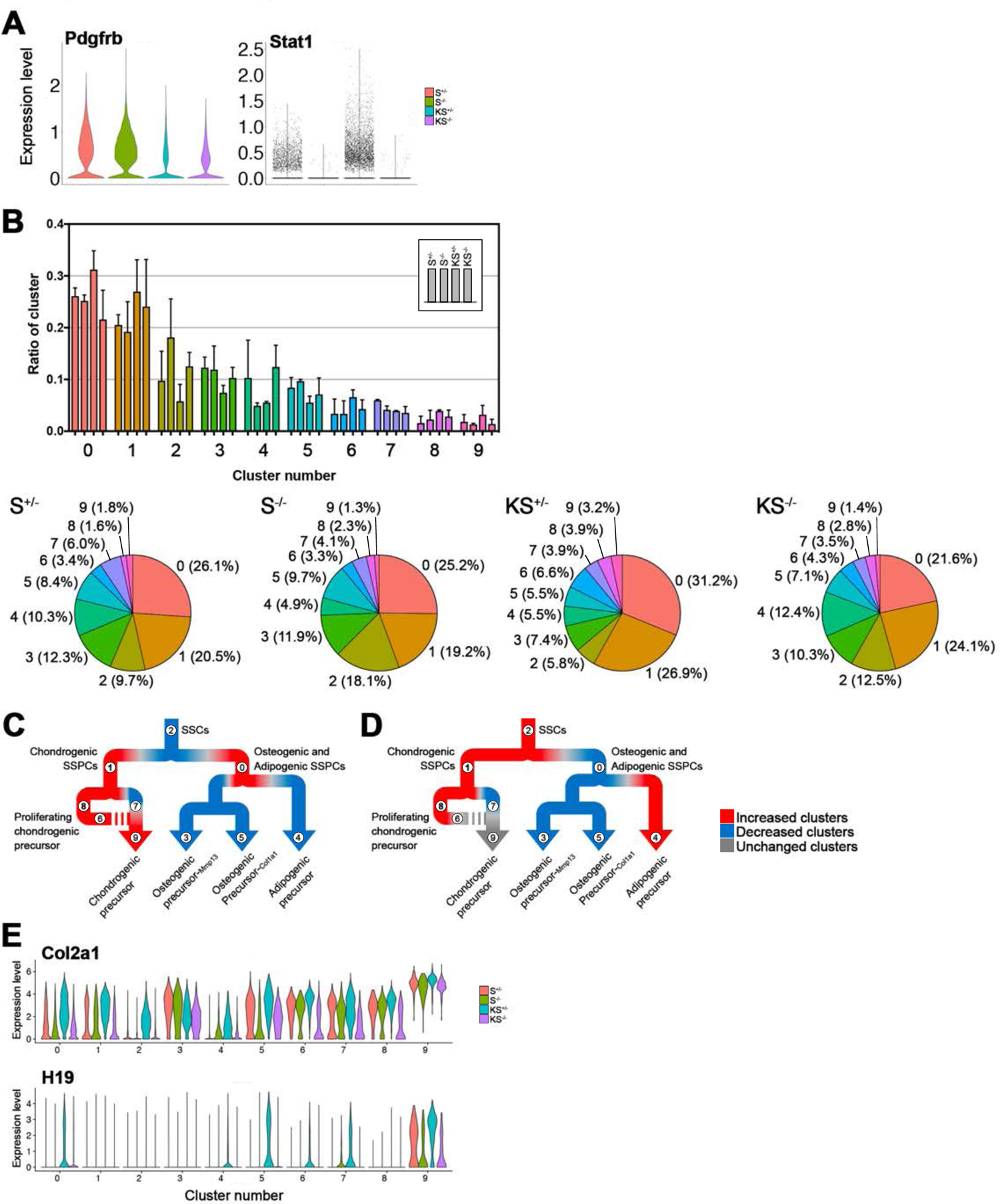
Single-cell RNA sequencing analysis reveals genotype-specific clusters and gene expression. **(A)** scRNA data were subgrouped by four genotypes. Expression levels of *Pdgfrb* and *Stat1* between genotypes. **(B)** Proportional ratio of each cluster in four genotypes were plotted (above) and also presented as pie charts with ratio labelling (below). **(C-D)** Color-coded *KS^+/-^* mutant (C) and *KS^-/-^* mutant (D) with their increase (red), decrease (blue) and no change status (gray) based on Supplementary Figure 5B’s ratio. **(E)** Differential expression of *Col2a1* and *H19* throughout 10 clusters between four genotypes.

**Supplementary Figure 6.**
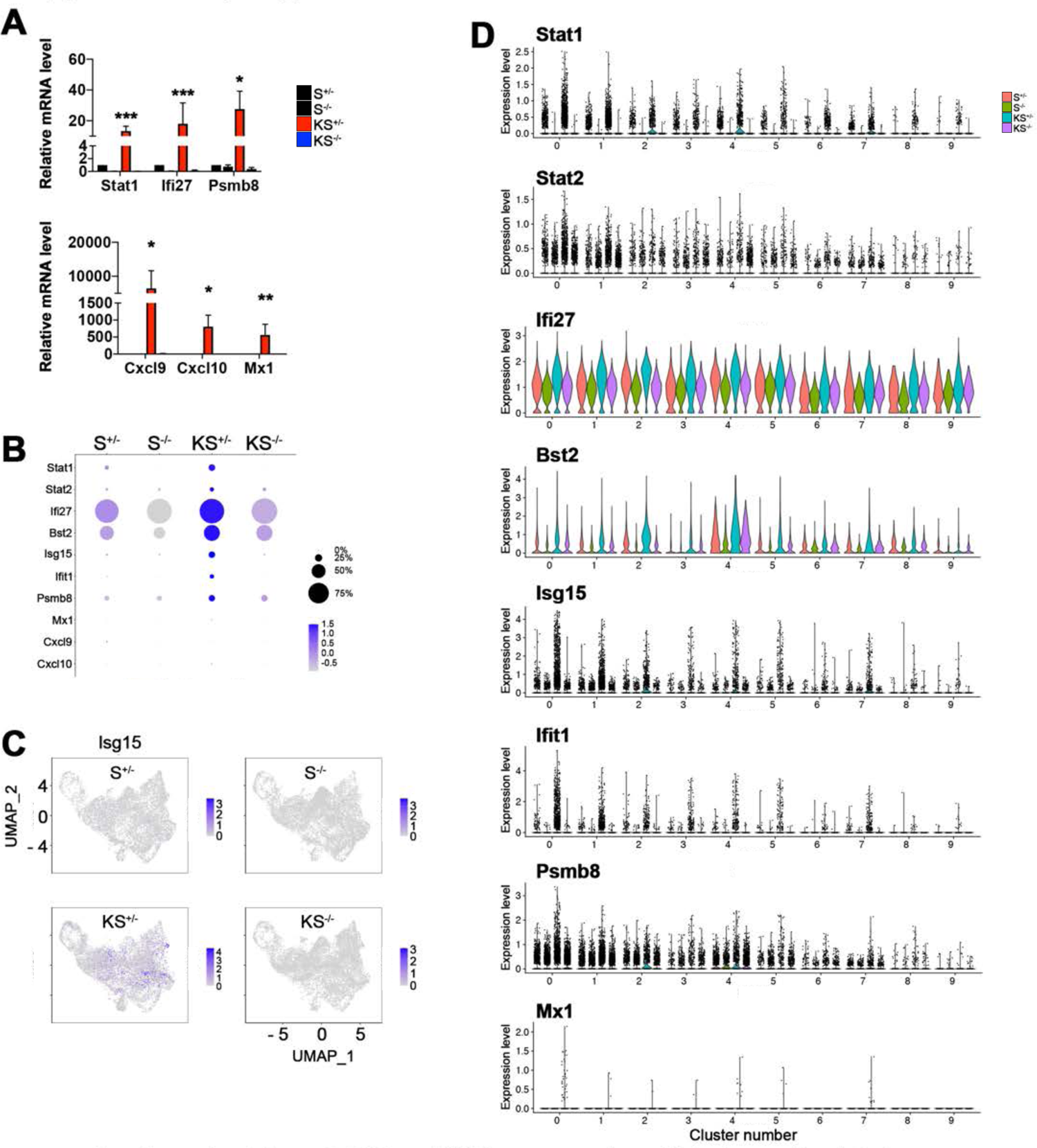
SSCs exhibit low expression of interferon-stimulated genes (ISGs). **(A)** Fold change in ISG mRNA levels in primary SSPCs isolated from 3-week-old limbs, by qRT-PCR with normalization to *Gapdh* and *Timm17b* (n = 5 per genotype). **(B)** Expression levels of *Stat1*, *Stat2* and ISGs between four genotypes in scRNA colonies. **(C)** UMAP plots to visualize distribution of *Isg15*-positive cells throughout 10 clusters. **(D)** Differential expression levels of *Stat1*, *Stat2* and ISGs (*Ifi27*, *Bst2*, *Isg15*, *Ifit1*, *Psmb8* and *Mx1*) throughout 10 clusters between four genotypes. ***, *p* < 0.005, **, *p* < 0.01 and *, *p* < 0.05 by one-way ANOVA. Data represent mean ± SEM.

**Supplementary Table 1.** Cluster markers for plotting the heatmap in Figure 3B.

| **Cluster #** | **Gene markers** |
| --- | --- |
| Cluster 0 | Penk, 1500015O10Rik, Col3a1, Cdh11, C4b, Cxcl12, Clu, Mme, Ogn, Igf1 |
| Cluster 1 | S100a4, Timp3, Lgals3, S100a6, Ly6a, Dlk1, Ccnd1, Vim, S100a10, Sh3bgrl3 |
| Cluster 2 | Lcn2, Mmp3, Saa3, Thbs4, Ccl2, Ccl7, Cxcl5, Mt2, Dcn, Mt1 |
| Cluster 3 | Mmp13, Cxcl14, Thbs1, Cdkn1a, Fhl2, Fgf7, Hist1h2bc, Ibsp, Igfbp5 |
| Cluster 4 | Lpl, Fabp4, Car3, Hp, Rbp4, Adipoq, Wfdc21, G0s2, Apoe, Mgst1 |
| Cluster 5 | Acta2, Tagln, Myl9, Col1a1, Nupr1, Col1a2, Tpm1, Cnn1 |
| Cluster 6 | Ube2c, Hist1h1b, Birc5, Pclaf, Hmgb2, H2afz, Top2a, Stmn1, Cenpf, Prc1 |
| Cluster 7 | Prelp, Fn1, Adamts5, Ly6c1, Lox, Sema3c |
| Cluster 8 | Cenpa, Selenoh, Cdkn3, Ran, Ccnb2, Tubb4b |
| Cluster 9 | Col2a1, Ucma, Cnmd, Col11a1, 3110079O15Rik, Hapln1, Peg3, Acan, Col9a1, Cdkn1c |

**Supplementary Table 2.** GO and KEGG pathway annotation of signature gene markers in 10 clusters.

| Cluster # | GO and KEGG pathway (top5) | Genes in the GO and KEGG pathway | p-Value |
| --- | --- | --- | --- |
| Cluster 0 | Secreted | Htra1, 1500015O10Rik, Smoc1, Adamts1, Comp, Cp, Cxcl12, Col3a1, Col12a1, Col14a1, Crisp1d1, Esm1, Fbln1, Fbln7, Gas6, Gdf10, Islr, Igf1, Lum, Ogn, Omd, Pi15, Penk, Prss23, Sfrp4, Spp1, Srgn, C4b, Clu, Tgfb2 | 2.5E-20 |
|  | Extracellular region part | Abi3bp, Smoc1, Adamts1, Comp, Cp, Cxcl12, Col3a1, Col12a1, Col14a1, Fbln1, Fbln7, Gdf10, Igf1, Lum, Ogn, Omd, Spp1, Srgn, C4b, Tgfb2 | 2.9E-12 |
|  | Heparin binding | Abi3bp, Adamts1, Comp, Fgfr2, Fbln7, Thbs2 | 4.1E-6 |
|  | ECM organization | Abi3bp, Smoc1, Col3a1, Fbln1, Pdgfra, Tgfb2 | 9.7E-6 |
|  | Growth factor | Cxcl12, Gdf10, Igf1, Ogn, Tgfb2 | 3.7E-4 |
| Cluster 1 | Secreted | Capg, Cdsn, Ecm1, Fn1, Fbln2, Il1rl1, Ltbp1, Lgals1, Mgp, Prg4, Slurp1, Sema3c, Serpine2, Sbsn, Timp1, Timp3, Tinagl1 | 4.6E-7 |
|  | Extracellular region | Capg, Cdsn, Ecm1, Fn1, Fbln2, Il1rl1, Ltbp1, Lgals1, Lgals3, Mgp, Prg4, Slurp1, Sema3c, Serpine2, Sbsn, Timp1, Timp3, Tinagl1 | 3.8E-6 |
|  | CD59 antigen | Ly6a, Ly6c1, Slurp1 | 2.4E-3 |
|  | Carbohydrate binding | Fn1, Lgals1, Lgals3, Prg4, Serpine2, Tinagl1 | 2.0E-3 |
|  | Enzyme inhibitor activity | Anxa1, Serpinb6a, Serpine2, Anxa3, Timp1, Timp3 | 6.0E-4 |
| Cluster 2 | Secreted | 1500015O10Rik, Wisp2, Adm, Comp, Chl1, Ccl2, Ccl7, Cxcl1, Cxcl12, Col1a2, Col3a1, Col12a1, C3, Dcn, Ecm1, Fbln1, Fbln2, Gas6, Igf1, Igfbp3, Igfbp6, Igfbp7, Ltbp2, Lcn2, Lbp, Lox, Mmp3, Nbl1, Ptx3, Prelp, Rnase4, Sfrp2, Spp1, Slpi, Serpina3n, Serpine2, Serpinf1, Saa3, Cxcl5, Clu, Sod3, Timp1, Timp2 | 1.3E-30 |
|  | Extracellular region part | Abi3bp, Comp, Chl1, Ccl2, Ccl7, Cxcl1, Cxcl12, Col1a2, Col3a1, Col12a1, C3, Dcn, Ecm1, Fbln1, Fbln2, Igf1, Igfbp3, Lbp, Lox, Mmp3, Prelp, Spp1, | 7.5E-19 |
|  |  | Serpine2, Serpinf1, Saa3, Cxcl5, Sod3, Timp1, Timp2 |  |
|  | Growth factor binding | Wisp2, Col1a2, Col3a1, Igfbp3, Igfbp6, Igfbp7, Il6st, Ltbp2, Pdgfra | 2.2E-10 |
|  | EGF-like calcium binding | Comp, Fbln1, Fbln2, Gas6, Ltbp2, Lrp1, Sned1, Thbs4 | 4.0E-8 |
|  | Cell adhesion | Wisp2, Comp, Chl1, Col12a1, Igfbp7, Spp1, Thbs4, Vcam1 | 4.5E-4 |
| Cluster 3 | Secreted | Wif1, Bglap2, Cp, Cxcl1, Cxcl14, Cx3cl1, Ctgf, Dcn, Dkk3, Esm1, Efemp1, Fgf7, Fmod, Fn1, Fst, Igfbp5, Kitl, Lum, Loxl2, Mmp13, Ptx3, Pltp, Ibsp, Cxcl5, Adamts5, Tnc, Tnfrsf11b, Wnt5a | 1.9E-13 |
|  | Extracellular region part | Cp, Cxcl1, Cxcl14, Cx3cl1, Ctgf, Dcn, Dkk3, Fmod, Fn1, Igfbp5, Kitl, Lum, Loxl2, Mmp13, Cxcl5, Adamts5, Tnc, Thbs1, Tnfrsf11b, Wnt5a | 6.0E-10 |
|  | Ossification | Jund, Ctgf, Fhl2, Igfbp5, Mmp13, Ptgs2, Shox2, Ibsp | 5.0E-7 |
|  | Chemokine (IL8-like) | Cxcl1, Cxcl14, Cx3cl1, Cxcl5 | 3.8E-4 |
|  | IGFBP | Ctgf, Esm1, Igfbp5 | 2.4E-3 |
| Cluster 4 | Secreted | S100a13, Smoc1, Adipoq, Angpt1, Agt, Apoe, Comp, Ccl9, Cxcl12, Col4a1, Col4a2, Cfd, Esm1, Efemp1, Fbln5, Fstl1, Gsn, Gpx3, Gas6, Ghr, Hp, Islr, Igf1, Igfbp7, Il1rn, Kng1, Kitl, Pi15, Retn, Rarres2, Rbp4, Sfrp1, Srgn, Serping1, C4b, Nid1, Adamts5, Stc1, Sbsn, Trf, Vcan | 2.9E-14 |
|  | Triglyceride metabolic process | Abhd5, Cat, Cav1, Dgat1, Gpx1, Lpl, Pnpla2 | 3.3E-7 |
|  | Lipid droplet | Abhd5, Pnpla2, Plin1, Plin4 | 4.4E-5 |
|  | Regulation of lipid metabolic process | Abhd5, Adipoq, Agt, Apoe, Cav1, Irs1, Pnpla2 | 6.0E-6 |
|  | Sulfur metabolic process | Cdo1, Ggt5, Gpx3, Ghr, Igf1, Idh1, Mgst1 | 9.6E-5 |
| Cluster 5 | Focal adhesion | Actb, Actg1, Actn1, Col1a1, Col1a2, Col5a2, Col11a1, Flna, Ibsp, Myl9, Thbs1, Vcl | 1.2E-8 |
|  | Extracellular matrix | Aspn, Bmp4, Ccdc80, Col1a1, Col1a2, Col5a2, Col11a1, Col12a1, Ctgf, Fmod, Npnt, Ogn, Omd | 3.1E-11 |
|  | Actin binding | Actn1, Cald1, Cnn1, Cnn2, Dstn, Flna, Myh10, Myh9, Palld, Tmsb4x, Tagln, Tpm1, Tpm2, Vcl | 2.1E-12 |
|  | Skeletal system development | Bmp4, Cdh11, Col1a1, Col1a2, Col5a2, Col11a1, Col12a1, Ctgf, Fhl2, Igfbp4, Igfbp5, Ibsp, Tnfrsf11b | 1.0E-8 |
|  | Contractile fiber part | Acta2, Actg2, Actn1, Fhl2, Hspb1, Myl9, Tpm1, Tpm2, Vcl | 3.9E-8 |
| Cluster 6 | Chromosome | Asf1b, Dnmt1, Prim1, H1f0, H1fx, H2afx, H2afz, Ndc80, Rad18, Rad21, Rad51, Rangap1, Ska1, Hjurp, Spc24, Spc25, Mki67, Aurkb, Birc5, Banf1, Bub1, Bub1b, Cdca8, Cenpa, Cenpe, Cenpf, Cenph, Cenpk, Cenpm, Cenpp, Cenpq, Clspn, Ercc6l, Hells, Hnrnpd, Hmga1, Hmgn2, Hist1h1a, Hist1h1b, Hist1h1d, Hist1h1e, Hist1h2ae, Hist1h4d, Hist2h2ac, Incenp, Kif22, Mcm2, Ncapd2, Ncapd3, Hmgb3, Hmgb1, Hmgb2, Hmga2, Pcna, Rfc2, Rfc4, Rfc5, Rpa2, Nuf2, Mad2l1, Smc2, Smc4, Tmpo, Tipin, Top2a, Topbp1 | 3.9E-49 |
|  | Cell cycle | Dbf4, E2f8, Fbxo5, H2afx, Ndc80, Rad21, Rad51, Ranbp1, Ran, Ska1, Hjurp, Racgap1, S100a6, Spc24, Spc25, Tpx2, Anp32b, Anln, Anxa1, Mki67, Aspm, Aurka, Aurkb, Birc5, Brca2, Bub1, Bub1b, Bub3, Cdk1, Cdc20, Cdc45, Cdca2, Cdca3, Cdca8, Cgref1, Cenpe, Cenpf, Cenph, Cep55, Chaf1a, Chaf1b, Clspn, Ccna2, Ccnb2, Ccnd1, Ccne1, Ccne2, Cdkn2d, Cdkn3, Ckap2, Ckap5, Dlgap5, Esco2, Ercc6l, Fam83d, Gmnn, Hells, Incenp, Kif11, Kif20b, Lig1, Mns1, Mcm2, Mcm3, Mcm6, Mcm7, Ncapd2, Ncaph, Ncapd3, Ncapg2, Nasp, Nusap1, Pdpn, Plk1, Cks1b, Phgdh, Calm1, Hmga2, Ccnb1, Psmc3ip, Prc1, Cks2, Nuf2, Gas2l3, Mad2l1, Stmn1, Smc2, Smc4, Trip13, Tipin, Topbp1, Tacc3, Tubb5, Ube2c, Uhrf1 | 6.2E-63 |
|  | DNA metabolic process | Dbf4, Dnmt1, Prim1, Gins2, H2afx, Rad18, Rad21, Rad51ap1, Rad51, Banf1, Brca2, Cdc45, Chaf1a, Chaf1b, Clspn, Ccne1, Ccne2, Dtl, Esco2, Fen1, Hells, Kif22, Lig1, Mcm2, Mcm3, Mcm4, Mcm5, Mcm6, Mcm7, Nasp, Hmgb2, Tk1, Pcna, Psmc3ip, Rfc2, Rfc3, Rfc4, Rfc5, Rpa2, Rrm1, Rrm2, Trip13, Tipin, Top2a, Topbp1, Usp1, Uhrf1 | 3.2E-23 |
|  | Ubl conjugation | Dnmt1, H2afx, H2afz, Rangap1, Ran, Anln, Aurka, Birc5, Bub1, Bub1b, Bub3, Cdk1, Cdc20, Cdca3, | 7.4E-16 |
|  |  | Cdca8, Cenpe, Ccnd1, Dlgap5, Hnrnpa1, Hist1h2ae, Hist1h4d, Hist2h2ac, Id1, Kif22, Lig1, Nusap1, Plk4, Hmgb1, Ube2s, Tuba1b, Calm1, Ccnb1, Set, Pcna, Nme1, Topbp1, Tubb5, Usp1, Ube2c, Uhrf1 |  |
|  | Microtubule cytoskeleton | Fbxo5, Ranbp1, Ska1, Racgap1, Tpx2, Aspm, Aurka, Aurkb, Birc5, Cdca8, Cenpe, Cenpf, Cep55, Ckap2, Ckap5, Dlgap5, Fam83d, Incenp, Kif11, Kif15, Kif20a, Kif20b, Kif22, Kif23, Kif4, Nusap1, Plk4, Tuba1b, Calm1, Ccnb1, Prc1, Kif2c, Nme1, Mad2l1, Stmn1, Tacc3, Tuba1c, Tubb5, Tubb6 | 3.5E-17 |
| Cluster 7 | Extracellular region part | Csf1, Fmod, Fn1, Fbln2, Grem1, Igfbp3, Lox, Masp1, Mfap5, Prelp, Sema3c, Adamts5, Sod3, Thbd, Thbs1, Timp2, Timp3 | 5.3E-11 |
|  | Calcium binding | S100a13, S100a6, Fbln2, S100a4, Thbs1 | 4.8E-6 |
|  | EGF-like region | Fn1, Fbln2, Ltbp2, Masp1, Thbd, Thbs1 | 9.7E-4 |
|  | Cell-substrate adhesion | Csf1, Fbln2, Thbs1 | 2.3E-3 |
|  | p53 signaling pathway | Fn1, Fbln2, Ltbp2, Masp1, Thbd, Thbs1 | 1.8E-5 |
| Cluster 8 | Cell cycle | H2afx, Rad21, Ranbp1, Ran, Racgap1, S100a6, Spc24, Tpx2, Anp32b, Anln, Anxa1, Mki67, Birc5, Cdc20, Cdca3, Cdca8, Cgref1, Cenpe, Cenpf, Ccna2, Ccnb2, Ccnd1, Cdkn3, Ckap2, Dlgap5, Nasp, Pdpn, Cks1b, Phgdh, Calm1, Hmga2, Ccnb1, Cks2, Nuf2, Mad2l1, Stmn1, Smc2, Tacc3, Tubb5, Ube2c | 1.0E-21 |
|  | Cytoskeleton | H1fx, H2afx, H2afz, Lmo1, Lyar, Nhp2, Nop58, Rad21, Ranbp1, Racgap1, Spc24, Tpx2, Actb, Anln, Mki67, Birc5, Banf1, Cdca8, Cenpa, Cenpe, Cenpf, Cenpm, Cenpp, Ckap2, Dlgap5, Dkc1, Eif6, Exosc8, Ezr, Hnrnpd, Hmga1, Hmgn2, Kif20a, Lmna, Msn, Ncl, Plec, Hmgb3, Hmgb1, Hmgb2, Tuba1b, Nop56, Calm1, Hmga2, Ccnb1, Actg1, Pa2g4, Nme1, Nuf2, Mad2l1, Stmn1, Smc2, Tmpo, Tacc3, Tpm2, Tubb5, Tubb6, Vim | 7.0E-16 |
|  | Chromosomal protein | H1fx, H2afx, H2afz, Birc5, Banf1, Cdca8, Cenpa, Cenpm, Cenpp, Hnrnpd, Hmga1, Hmgn2, Hmgb3, Hmgb1, Hmgb2, Hmga2, Tmpo | 9.2E-14 |
|  | Ubl conjugation | H2afx, H2afz, Ran, Anln, Birc5, Cdc20, Cdca3, Cdca8, Cenpe, Ccnd1, Dlgap5, Hnrnpa1, Id1, | 1.1E-8 |
|  |  | Hmgb1, Ube2s, Tuba1b, Calm1, Ccnb1, Set, Nme1, Tubb5, Ube2c |  |
|  | Macromolecular complex subunit | H1fx, H2afx, H2afz, Lsm2, Cav1, Cenpa, Dctpp1, Eif6, Nasp, Tuba1b, Set, Rrm1, Rrm2, Nap1l1, Stmn1, Tubb5, Tubb6 | 5.8E-7 |
| Cluster 9 | Extracellular region | Clec11a, Cmtm5, Edil3, 1500015O10Rik, S100b, Wfdc12, Acan, Cpe, Comp, Ccdc80, Col2a1, Col9a1, Col9a2, Col11a1, Col11a2, Col12a1, Col27a1, Ctgf, Cdsn, Crisp1d1, Dcn, Enpp2, Fmod, Fbln7, Fibin, Frzb, Grem1, Gdf10, Gdf15, Hspa5, Hapln1, 1190002N15Rik, Islr, Igfbp5, Il17d, Lum, Matn1, Matn4, Mgp, Msmp, Ogn, Omd, Pdgfrl, Pcolce2, Scrg1, Slpi, Serpina3n, Srp2x, Thbs1, Timp3, Tgfb, Ucn3 | 3.5E-22 |
|  | Secreted | Clec11a, Edil3, 1500015O10Rik, Wfdc12, Acan, Cpe, Comp, Ccdc80, Col2a1, Col9a1, Col9a2, Col11a1, Col11a2, Col12a1, Col27a1, Ctgf, Cdsn, Crisp1d1, Dcn, Enpp2, Fmod, Fbln7, Fibin, Frzb, Grem1, Gdf10, Gdf15, Hapln1, 1190002N15Rik, Islr, Igfbp5, Lum, Matn4, Mgp, Msmp, Ogn, Omd, Pdgfrl, Pcolce2, Scrg1, Slpi, Serpina3n, Srp2x, Timp3, Tgfb, Ucn3 | 2.3E-21 |
|  | Cartilage development | Sox9, Acan, Chst11, Col2a1, Col9a1, Col11a1, Col11a2, Ctgf, Cyl1, Fgfr3, Mgp, Pth1r | 3.1E-12 |
|  | Pattern binding | Acan, Comp, Ccdc80, Ctgf, Dcn, Enpp2, Fbln7, Hapln1, Pcolce2, Thbs1 | 7.8E-8 |
|  | Skeletal system development | Papss2, Sox9, Acan, Chst11, Col2a1, Col9a1, Col11a1, Col11a2, Ctgf, Cyl1, Fgfr3, Igfbp5, Mgp, Pth1r | 4.5E-8 |

**Supplementary Table 3.**
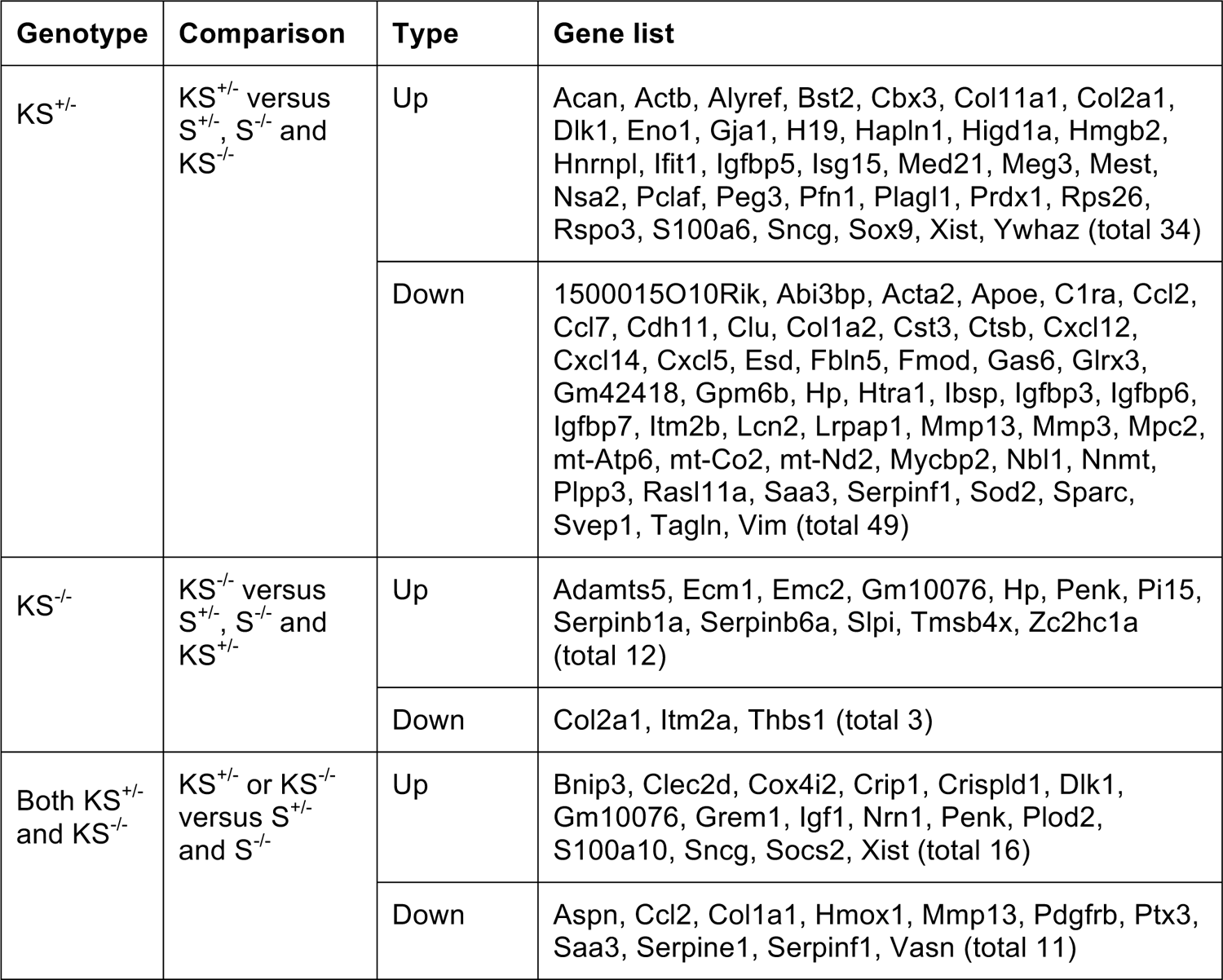
Upregulated or downregulated genes in cultured PDGFRβ GOF SSCs.

| Genotype | Comparison | Type | Gene list |
| --- | --- | --- | --- |
| KS<sup>+/-</sup> | KS<sup>+/-</sup> versus S<sup>+/-</sup>, S<sup>-/-</sup> and KS<sup>-/-</sup> | Up | Acan, Actb, Alyref, Bst2, Cbx3, Col11a1, Col2a1, Dlk1, Eno1, Gja1, H19, Hapln1, Higd1a, Hmgb2, Hnrnp1, Ifit1, Igfbp5, Isg15, Med21, Meg3, Mest, Nsa2, Pclaf, Peg3, Pfn1, Plagl1, Prdx1, Rps26, Rspo3, S100a6, Sncg, Sox9, Xist, Ywhaz (total 34) |
|  |  | Down | 1500015O10Rik, Abi3bp, Acta2, Apoe, C1ra, Ccl2, Ccl7, Cdh11, Clu, Col1a2, Cst3, Ctsb, Cxcl12, Cxcl14, Cxcl5, Esd, Fbln5, Fmod, Gas6, Glrx3, Gm42418, Gpm6b, Hp, Htra1, Ibsp, Igfbp3, Igfbp6, Igfbp7, Itm2b, Lcn2, Lrpap1, Mmp13, Mmp3, Mpc2, mt-Atp6, mt-Co2, mt-Nd2, Mycbp2, Nbl1, Nnmt, Plpp3, Rasl11a, Saa3, Serpinf1, Sod2, Sparc, Slep1, Tagln, Vim (total 49) |
| KS<sup>-/-</sup> | KS<sup>-/-</sup> versus S<sup>+/-</sup>, S<sup>-/-</sup> and KS<sup>+/-</sup> | Up | Adamts5, Ecm1, Emc2, Gm10076, Hp, Penk, Pi15, Serpinb1a, Serpinb6a, Slpi, Tmsb4x, Zc2hc1a (total 12) |
|  |  | Down | Col2a1, Itm2a, Thbs1 (total 3) |
| Both KS<sup>+/-</sup> and KS<sup>-/-</sup> | KS<sup>+/-</sup> or KS<sup>-/-</sup> versus S<sup>+/-</sup> and S<sup>-/-</sup> | Up | Bnip3, Clec2d, Cox4i2, Crip1, Crisp1d1, Dlk1, Gm10076, Grem1, Igf1, Nrn1, Penk, Plod2, S100a10, Sncg, Socs2, Xist (total 16) |
|  |  | Down | Aspn, Ccl2, Col1a1, Hmox1, Mmp13, Pdgrfb, Ptx3, Saa3, Serpine1, Serpinf1, Vasn (total 11) |

**Supplementary Table 4.** GO and KEGG annotation of upregulated or downregulated genes in cultured PDGFRβ GOF SSCs.

| Genotype | Type | GO/KEGG (Top5) | Genes in the GO and KEGG pathway | p-value |
| --- | --- | --- | --- | --- |
| KS<sup>+/-</sup> | Up<br>(among<br>total 34) | Chordate embryonic development | Col2a1, Col11a1, Dlk1, Med21, Pfn1 | 3.4E-3 |
|  |  | Cartilage development | Sox9, Acan, Col2a1, Col11a1 | 6.0E-6 |
|  |  | Regulation of apoptosis | Sox9, Col2a1, Prdx1, Ywhaz | 6.5E-2 |
|  |  | Glycosaminoglycan binding | Rspo3, Acan, Hapln1 | 2.3E-2 |
|  |  | Regulation of transcription | Sox9, Eno1, Hmgb2, Med21, Plagl1, Pfn1 | 1.1E-2 |
|  | Down<br>(among<br>total 49) | Secreted factors | Htra1, 1500015O10Rik, Apoe, Ccl2, Ccl7, Cxcl12, Cxcl14, Col1a2, Cst3, Fmod, Fbln5, Gas6, Hp, Igfbp3, Igfbp6, Igfbp7, Lcn2, Mmp13, Mmp3, Nbl1, Sparc, Serpinf1, Saa3, Ibsp, Cxcl5, Clu, Svep1 | 3.9E-19 |
|  |  | Growth factor binding | Htra1, Igfbp3, Igfbp6, Igfbp7 | 5.0E-6 |
|  |  | Chemokine | Ccl2, Ccl7, Cxcl12, Cxcl14, Cxcl5 | 1.5E-6 |
|  |  | Extracellular matrix | Col1a2, Fmod, Mmp13, Mmp3, Sparc | 1.4E-3 |
|  |  | Heparin binding | Abi3bp, Apoe, Ccl7, Lrpap1 | 1.1E-3 |
| KS<sup>-/-</sup> | Up<br>(among<br>total 12) | Protease inhibitor | Pi15, Slpi, Serpinb1a, Serpinb6a | 1.2E-5 |
|  |  | Extracellular region | Ecm1, Hp, Pi15, Penk, Slpi, Adamts5, Tmsb4x | 5.8E-6 |
|  | Down<br>(among<br>total 3) | ECM-receptor interaction | Col2a1, Thbs1 | 1.2E-2 |
|  |  | Focal adhesion | Col2a1, Thbs1 | 2.7E-2 |
| Both KS<sup>+/-</sup><br>and KS<sup>-/-</sup> | Up<br>(among<br>total 16) | Disulfide bond | Clec2d, Crispld1, Cox4i2, Dlk1, Grem1, Igf1, Penk | 6.9E-3 |
|  |  | Anti-apoptosis | Bnip3, Igf1, Socs2 | 9.6E-3 |
|  |  | Extracellular region | Crispld1, Dlk1, Grem1, Igf1, Penk | 5.8E-2 |
|  |  | Glycoprotein | Clec2d, Dlk1, Grem1, Nrn1, Plod2 | 3.3E-1 |
|  |  | Metal ion binding | S100a10, Crip1, Dlk1, Plod2 | 6.7E-1 |
|  | Down<br>(among<br>total 11) | Extracellular matrix | Aspn, Ccl2, Col1a1, Mmp13, Ptx3, Serpine1, Serpinf1, Saa3 | 1.6E-5 |
|  |  | Skeletal system development | Col1a1, Mmp13, Pdgfrb | 4.9E-3 |
|  |  | Extracellular space | Ccl2, Serpinf1, Saa3 | 4.4E-2 |
|  |  | Disulfide bond | Aspn, Ccl2, Mmp13, Ptx3 | 7.7E-1 |

**Supplementary Table 5.**
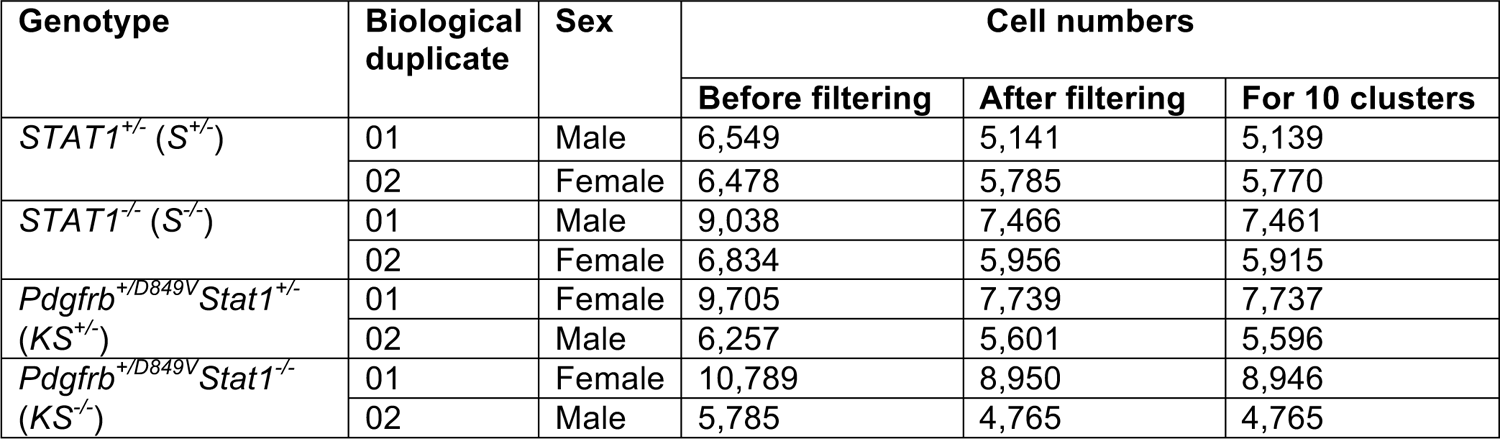
Genotypes and cell numbers for single-cell RNA sequencing.

| Genotype | Biological duplicate | Sex | Cell numbers |  |  |
| --- | --- | --- | --- | --- | --- |
|  |  |  | Before filtering | After filtering | For 10 clusters |
| *STAT1*<sup>+/-</sup> (*S*<sup>+/-</sup>) | 01 | Male | 6,549 | 5,141 | 5,139 |
|  | 02 | Female | 6,478 | 5,785 | 5,770 |
| *STAT1*<sup>-/-</sup> (*S*<sup>-/-</sup>) | 01 | Male | 9,038 | 7,466 | 7,461 |
|  | 02 | Female | 6,834 | 5,956 | 5,915 |
| *Pdgfrb*<sup>+/*D849V*</sup> *Stat1*<sup>+/-</sup> (*KS*<sup>+/-</sup>) | 01 | Female | 9,705 | 7,739 | 7,737 |
|  | 02 | Male | 6,257 | 5,601 | 5,596 |
| *Pdgfrb*<sup>+/*D849V*</sup> *Stat1*<sup>-/-</sup> (*KS*<sup>-/-</sup>) | 01 | Female | 10,789 | 8,950 | 8,946 |
|  | 02 | Male | 5,785 | 4,765 | 4,765 |

**Supplementary Table 6.** Cell numbers and ratios of individual clusters in biological duplicates in Supplementary Figure 5B.

| Genotype | *STAT1*<sup>+/-</sup> (*S*<sup>+/-</sup>) |  | *STAT1*<sup>-/-</sup> (*S*<sup>-/-</sup>) |  | *Pdgfrb*<sup>+D849V</sup>*Stat1*<sup>+/-</sup> (*KS*<sup>+/-</sup>) |  | *Pdgfrb*<sup>+D849V</sup>*Stat1*<sup>-/-</sup> (*KS*<sup>-/-</sup>) |  |
| --- | --- | --- | --- | --- | --- | --- | --- | --- |
| Duplicate | 01 | 02 | 01 | 02 | 01 | 02 | 01 | 02 |
| Sex | Male | Female | Male | Female | Female | Male | Female | Male |
| Cluster0 | 1421<br>(28%) | 1414<br>(25%) | 1963<br>(26%) | 1422<br>(24%) | 2696<br>(35%) | 1542<br>(28%) | 2435<br>(27%) | 760<br>(16%) |
| Cluster1 | 1156<br>(22%) | 1069<br>(19%) | 1866<br>(25%) | 792<br>(13%) | 1609<br>(21%) | 1852<br>(33%) | 1341<br>(15%) | 1580<br>(33%) |
| Cluster2 | 207<br>(4%) | 892<br>(15%) | 796<br>(11%) | 1511<br>(26%) | 196<br>(3%) | 506<br>(9%) | 882<br>(10%) | 725<br>(15%) |
| Cluster3 | 735<br>(14%) | 589<br>(10%) | 546<br>(7%) | 973<br>(16%) | 685<br>(9%) | 337<br>(6%) | 1102<br>(12%) | 393<br>(8%) |
| Cluster4 | 153<br>(3%) | 1014<br>(18%) | 326<br>(4%) | 323<br>(5%) | 411<br>(5%) | 322<br>(6%) | 1485<br>(17%) | 391<br>(8%) |
| Cluster5 | 533<br>(10%) | 371<br>(6%) | 747<br>(10%) | 552<br>(9%) | 522<br>(7%) | 241<br>(4%) | 921<br>(10%) | 186<br>(4%) |
| Cluster6 | 320<br>(6%) | 30<br>(1%) | 438<br>(6%) | 49<br>(1%) | 617<br>(8%) | 288<br>(5%) | 225<br>(3%) | 288<br>(6%) |
| Cluster7 | 300<br>(6%) | 354<br>(6%) | 363<br>(5%) | 200<br>(3%) | 297<br>(4%) | 224<br>(4%) | 207<br>(2%) | 227<br>(5%) |
| Cluster8 | 149<br>(3%) | 13<br>(0.2%) | 302<br>(4%) | 28<br>(0.5%) | 319<br>(4%) | 207<br>(4%) | 142<br>(2%) | 194<br>(4%) |
| Cluster9 | 165<br>(3%) | 24<br>(0.4%) | 114<br>(2%) | 65<br>(1%) | 385<br>(5%) | 77<br>(1%) | 206<br>(2%) | 21<br>(0.4%) |
| Total cells | 5139 | 5770 | 7461 | 5915 | 7737 | 5596 | 8946 | 4765 |

\*Numbers in parentheses indicate the percentages of individual clusters.

**Supplementary Table 7.**
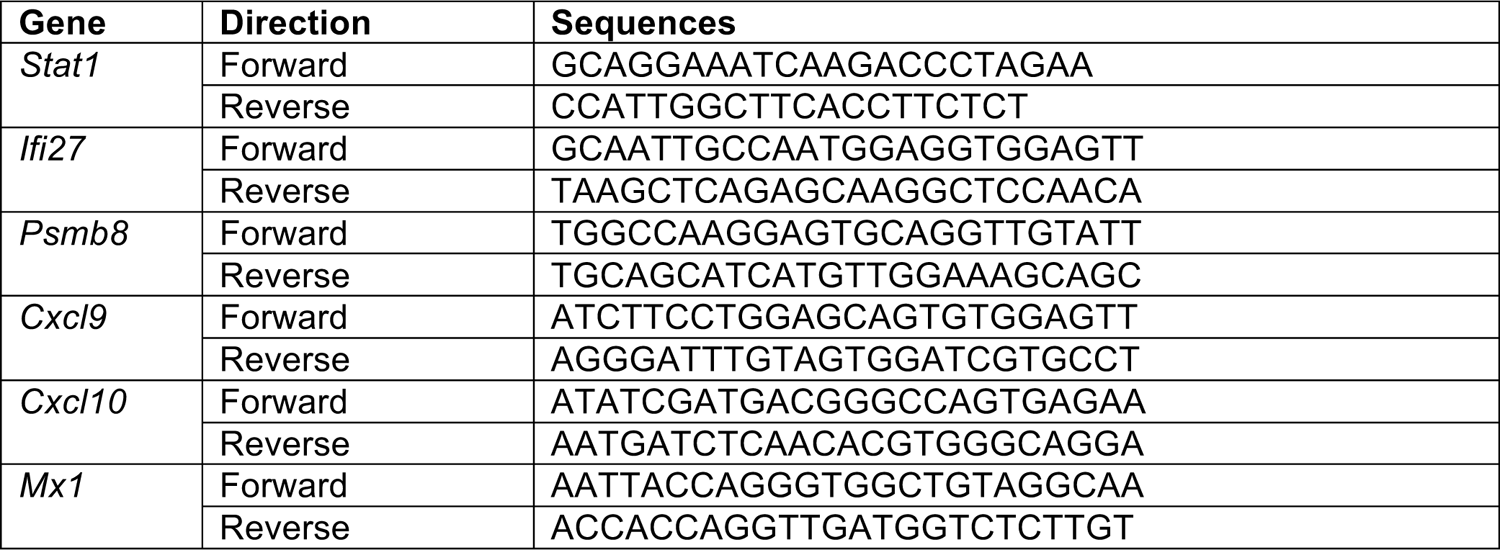
Primer sequences for RT-PCR.

**Supplementary Table 8.** Primary antibodies and fluorophore-conjugated antibodies used in this study.

| Use* | Epitope | Dilution | Source | Catalog# |
| --- | --- | --- | --- | --- |
| WB | STAT1 | 1:2000 | Millipore | 06-501 |
|  | STAT1 pY701 | 1:2000 | Invitrogen | 333400 |
|  | STAT5 | 1:2000 | Cell Signaling Technology | 94205 |
|  | STAT5 pY694 | 1:2000 | Cell Signaling Technology | 9351 |
|  | PDGFRb | 1:2000 | Santa Cruz | sc-432 |
|  | PDGFRb pY1009 | 1:2000 | Cell Signaling Technology | 3124 |
| IF | Sox9 | 1:500 | Millipore | AB5535 |
| FACS | CD31-PE/Cy7 | 1:2000 | BioLegend | 102417 |
|  | CD45-PE/Cy7 | 1:500 | BioLegend | 109829 |
|  | Sca1-APC/Cy7 | 1:100 | BioLegend | 108126 |
| | PDGFRα(CD140a)-APC | 1:20 | eBioscience | 17-1401-81 |
|  | TER-119-PE/Cy7 | 1:500 | BioLegend | 116221 |
|  | Zombie Green | 1:1000 | BioLegend | 423111 |

\*WB: western blotting
IF: immunofluorescence
FACS: fluorescence-activated cell sorting

